# Genetic Diversity of Cytochrome P450 Genes in *Apis mellifera* Subspecies

**DOI:** 10.64898/2026.03.20.713126

**Authors:** Fernanda Li, Daniela Lima, Sana Bashir, Carlos Yadró Garcia, Ana Rita Lopes, Gilles Verbinnen, Dirk C. de Graaf, Lina De Smet, Anais Rodriguez, Annelise Rosa-Fontana, José Rufino, Raquel Martín-Hernandez, Medibees Consortium, Maria Alice Pinto, Dora Henriques

## Abstract

The western honey bee (*Apis mellifera*) is an essential pollinator facing unprecedented threats from pesticide exposure. While pesticide resistance evolution is well documented in agricultural pests, our understanding of genetic variation in honey bee detoxification systems remains limited. This represents a missed opportunity, as harnessing naturally occurring detoxification diversity could provide new avenues for pollinator protection. Cytochrome P450 monooxygenases (CYPs), which are central to xenobiotic metabolism, offer a promising starting point. Here, we present the first comprehensive analysis of CYP genetic diversity in *A. mellifera*. We analysed the CYPome of 1,467 individuals representing 18 *A. mellifera* subspecies from 25 countries and identified 5,756 single-nucleotide polymorphisms (SNPs) in 46 CYP genes. Imputed McDonald-Kreitman testing revealed that 56% of non-synonymous CYP substitutions were driven by positive selection. Of the 1,302 haplotypes identified, 84% resided in CYP3, concentrated in the CYP9 and CYP6AS subfamilies implicated in xenobiotic detoxification. Population-level analysis of nucleotide diversity, Tajima’s D selection signatures, F_ST_-based differentiation, and McDonald-Kreitman testing pointed to CYP3 clan genes as the primary locus of adaptive variation.

This work provides the first step toward building a comprehensive pharmacogenomic resource for honey bees, enabling the prediction of population-specific pesticide vulnerabilities and leveraging naturally occurring detoxification variants to enhance pollinator resilience – a critical step toward sustainable pollinator management.

## 1. Introduction

The worldwide decline in diversity and number of pollinating species poses a major threat to both food security and ecosystem health (Goulson et al., 2015; Klein et al., 2007; Potts et al., 2010). Although this decline is multifactorial, pesticides are a primary concern; while designed to eliminate pest species, their effects seldom remain confined to the intended targets (Wan et al., 2025).

The western honey bee, *Apis mellifera L*., a crucial pollinator of crops worldwide, is among the non-target species most exposed to pesticides, with impacts observed across multiple levels, from individual physiology and behaviour to colony survival (reviewed in Pisa et al., 2017). Documented effects include acute mortality, impaired foraging and homing (e.g., with neonicotinoids), learning and memory deficits, neurotoxic symptoms, and abnormal behaviour (Arena and Sgolastra, 2014; Tosi et al., 2022). Because of this vulnerability and ecological importance, *A. mellifera* is the standard test species in pesticide risk assessments worldwide, including in the EU and the US (EFSA et al., 2023; OECD, 2017). Yet numerous studies report variation in median lethal dose (LD₅₀) values and other toxicity endpoints for individuals exposed to the same pesticide (Aupinel et al., 2010; Blacquière et al., 2012; Iwasa et al., 2004; Laurino et al., 2013). Evidence also shows clear differences among *A. mellifera* subspecies and genetic stocks (*A. m. ligustica*, *A. m. carnica*, *A. m. mellifera*, *A. m. caucasia,* as well as Russian and Africanised honey bees) in their sensitivity to identical compounds (Danka et al., 1986; Elzen et al., 2003; Gregorc et al., 2016; Iwasa et al., 2004; Laurino et al., 2013; Milone et al., 2020; Rinkevich et al., 2015; Suchail et al., 2000). *A. m. ligustica*, for instance, can be up to 34 times more sensitive to imidacloprid than *A. m. carnica* (Rinkevich et al., 2015), while Africanized bees show greater tolerance to azinphosmethyl, methyl parathion, and permethrin, but increased sensitivity to carbaryl (Danka et al., 1986; Rinkevich et al., 2015). Such patterns strongly suggest that, as in human pharmacogenomics, genetic variation is a key determinant of pesticide sensitivity in honey bees. Supporting this, the heritability of clothianidin resistance was recently estimated at 37.8% (Tsvetkov et al., 2023). Despite this experimental evidence for subspecific differences in pesticide sensitivity, the genetic variability of important pesticide response genes in *A. mellifera* remains poorly characterized, even though the species comprises 31 recognized subspecies grouped into four major evolutionary lineages: M (Western and Northeastern Europe, Northwestern China), C (Central and Southeastern Europe), A (African), and O (Near East and Central Asia).

Understanding the genetic mechanisms underlying pesticide tolerance in *A. mellifera* is still in its infancy. In other insect species, pesticide resistance has generally been linked to two processes: (1) target site insensitivity, in which point mutations make insect proteins targeted by xenobiotics insensitive, and (2) metabolic resistance, through an increase in the metabolic capabilities of detoxification enzymes. Because pesticides target essential insect proteins, only limited point mutations can reduce pesticide sensitivity without compromising function. In contrast, genes encoding detoxification enzymes exhibit greater plasticity, accumulating changes that alter gene expression and protein function. This plasticity is essential for these genes, as they mediate environmental responses and contribute to pesticide survival (reviewed in Nauen et al., 2022). Amplification, overexpression (non coding sequence variation) and coding sequence variation (altering enzyme function) are the primary mechanisms driving the increase in metabolic action against pesticides.

Cytochrome P450 monooxygenases (CYPs), commonly referred to as P450s, are central players in environmental response actions and the focus of our studys. They Clique ou toque aqui para introduzir texto.are essential in insects for detoxifying a broad range of exogenous compounds, including plant secondary metabolites, pesticides, and environmental pollutants (reviewed by Nauen et al., 2022). CYPs also metabolise endogenous compounds, such as ecdysteroids, juvenile hormone, cuticular hydrocarbons, pheromones, and other semiochemicals involved in physiological functions (Claudianos et al., 2006). According to their phylogenetic relationships, insect CYP genes are classified into six major clans: CYP2, CYP3, CYP4, CYP16, CYP20 and mitochondrial (mito) (Claudianos et al., 2006; Dermauw et al., 2020; Feyereisen, 2012). The size of the CYP gene repertoire, or CYPome, varies greatly among insects. For example, the body louse *Pediculus humanus* has only 36 CYPs, whereas the mosquito *Culex quinquefasciatus* has 196 (Dermauw et al., 2020). The western honey bee, *Apis mellifera,* sits at the lower end of this spectrum, with only 46 genes and 4 clans (Claudianos et al., 2006). This reduction is particularly marked in the CYP4 clan (only four genes) and by the complete absence of CYP16 and CYP20 clan, present only in certain species in Apterygota and Paleoptera (Dermauw et al., 2020), and mitochondrial CYP12 genes which have been linked to insecticide resistance in other species (Bogwitz et al., 2005; Claudianos et al., 2006; Daborn et al., 2007; Guzov et al., 1998; Shi et al., 2022). Despite this, the western honey bee is not universally more sensitive to pesticides. For example, *A. mellifera* (46 CYP genes) is less sensitive to N-cyanoamidine neonicotinoids than the leafcutter bee *Megachile rotundata* (49 CYP genes), a difference attributed to the absence of the CYP9Q subfamily in the latter (Haas et al., 2022b; Hayward et al., 2019; Manjon et al., 2018). Pesticide tolerance is ultimately determined by the presence of functionally relevant subfamilies, not the number of CYP genes.

In *A. mellifera*, both CYP6 and CYP9 (CYP 3 clan) families are involved in the direct detoxification of xenobiotics. However, while CYP6 is associated with the detoxification of plant allelochemichals, such as quercetin, CYP9, in particular, the CYP9Q subfamily (CYP9Q1, CYP9Q2, and CYP9Q3 genes), is central for pesticide detoxification (Mao et al., 2011). These enzymes metabolise compounds from at least three insecticide classes: neonicotinoids, pyrethroids, and organophosphates (Hayward et al., 2019; Manjon et al., 2018; Mao et al., 2011). Their function parallels that of human CYP3A4 and CYP2D6, which together metabolise a large fraction of clinically used drugs (Haas et al., 2022b; Zanger and Schwab, 2013). Both human genes are highly polymorphic, with genetic variation strongly influencing individual metabolic profiles and drug responses (Zanger and Schwab, 2013). In honey bees, single nucleotide polymorphisms (SNPs) in CYP9Q1 and CYP9Q3 coding sequences, have been associated with clothianidin tolerance, likely by affecting substrate recognition and metabolism (Tsvetkov et al., 2023). Nonetheless, the genetic diversity of these genes across honey bee subspecies and populations remains poorly characterised. In addition, because the function and substrate specificity of most honey bee P450s are unknown, documenting variation in these genes could highlight which CYPs are less amenable to sequence variation an, thus,s less likely to participate in xenobiotic detoxification. This also provides a pool of candidate variants that can be assessed for their impact on gene expression or protein activity in follow-up studies.

Here, we present an unprecedented large-scale analysis of genetic diversity across 46 CYP genes in *A. mellifera*, extracted from whole-genome sequencing data of 18 subspecies sampled across 25 countries. As genetic diversity encompasses a broad spectrum of molecular changes, we focused on SNPs to address the following questions: (1) Do certain CYP clans accumulate more polymorphisms than others, thus reflecting differences in the roles they play (endogenous vs. xenobiotic metabolism)? (2) Do CYP genes with essential metabolic roles show lower diversity than those involved in environmental responses, and are the most deleterious mutations concentrated in the latter? (3) Are the xenobiotic detoxification subfamilies enriched for signatures of positive selection? (4) Is there sufficient subspecies and country specific variation to support local adaptation to pesticide exposure? By answering these questions, we will gain a clearer picture of the selective pressures acting on each CYP gene and its role in xenobiotic metabolism, and identify which genes are most likely to process novel pesticides. This will reveal the extent to which natural variation in the *A. mellifera* CYPome can be leveraged to improve pesticide risk assessment.

## 2. Materials and methods

### 2.1 Dataset composition

In this study, we analysed the 46 CYP genes of 1,467 *A. mellifera* drones (haploid males) collected from 25 countries (Figures 1A and B) and representing 18 subspecies belonging to the four main evolutionary lineages (A, C, M, and O) (Figures 1C and D; Table S1). Most of the samples were collected from countries in the Mediterranean basin and the Middle East (e.g.,Algeria, Cyprus, Egypt, Greece, Iran, Italy, Jordan, Lebanon, Morocco, Oman,Turkey, Portugal, Spain, and the UAE), with additional samples from Europe (Croatia, Denmark, France, Malta, Netherlands, Norway, Serbia, Slovenia, Switzerland, and the UK), as well as Cuba. This geographic breadth captures a considerable portion of *A. mellifera* subspecific diversity as well as a wide range of floral diversity and agricultural practices, reflecting variation in honey bee exposure to plant secondary metabolites and pesticides.

**Figure 1.**
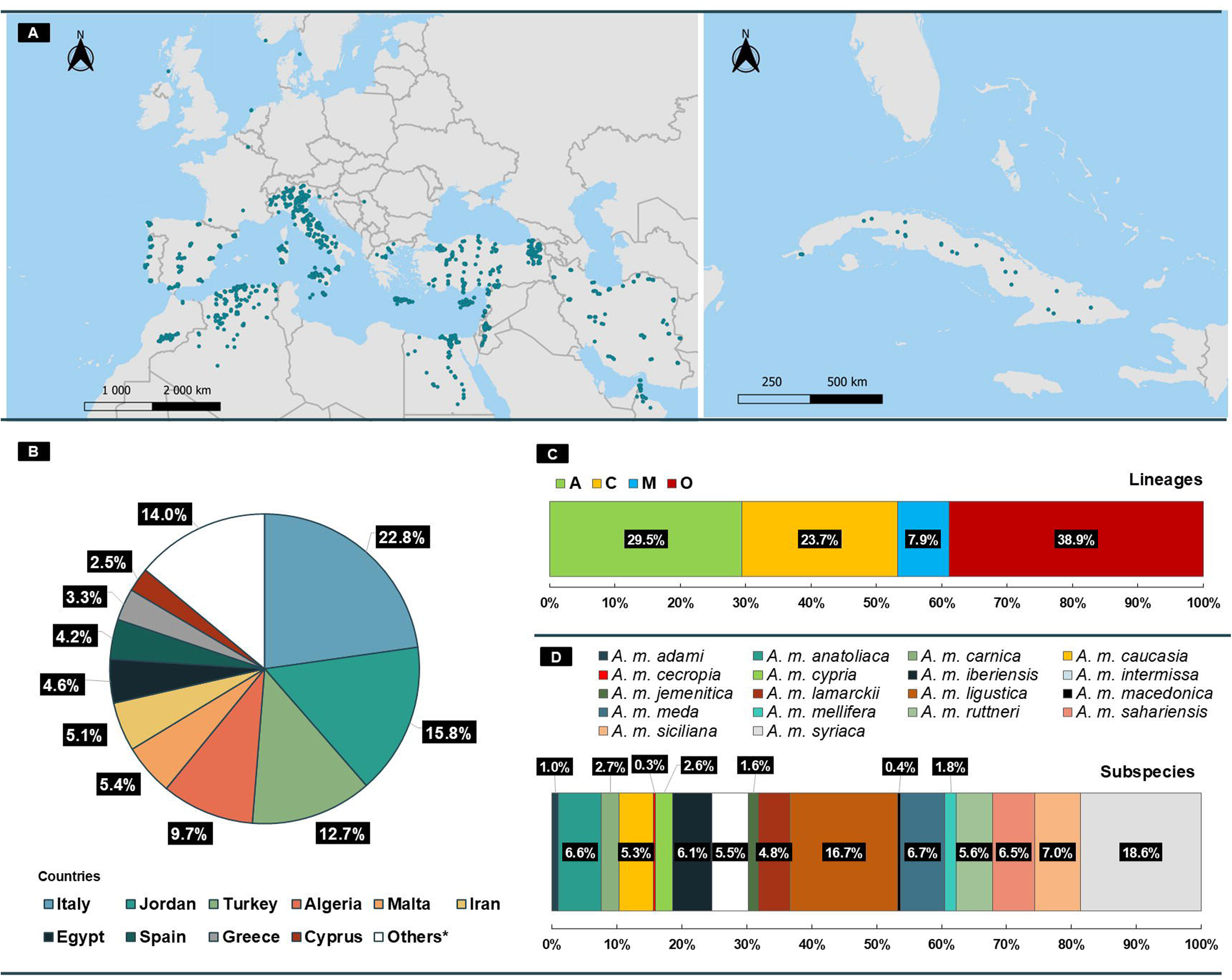
Haploid Dataset. Data were collected from 1,467 adult honey bees across 25 countries and 18 subspecies. **A –** Sample distribution. **B –** Proportion of samples per country. *This category includes the following 15 countries: Cuba (2.2%), Morocco (2.1%), Lebanon (2.0%), Portugal (1.7%), Switzerland (1.5%), France (1.4%), Oman (0.9%), Slovenia (0.9%), United Arab Emirates (0.7%), and Croatia, the Netherlands, Denmark, Norway, Serbia, and the United Kingdom (each 0.1%). **C –** Lineage of sampled honey bees. African (A) and Middle Eastern (O) lineages together account for 68.4% of the samples. **D –** Relative frequencies of honey bee subspecies. A. m. syriaca (18.6%) and A. m. ligustica (16.7%) are the most represented subspecies.

The CYPome data were extracted from whole-genome sequences generated within the framework of the MEDIBEES project, as well as from publicly available whole-genome sequences in the Sequence Read Archive (SRA, https://www.ncbi.nlm.nih.gov/sra), including PRJEB16533, PRJNA311274 (Wragg et al., 2022),PRJNA1305528, PRJNA1305509, and PRJNA1307299 (Yadro-Garcia et al., 2026). Sequencing and SNP calling approaches are detailed in Wragg et al. (2022) and Yadro-Garcia et al. (2026).

### 2.2 Sequencing and bioinformatic processing

All genomes were sequenced using Illumina technology (Illumina Inc., San Diego, CA, USA), albeit with different versions of the Illumina HiSeq platform, as detailed in Henriques et al. (2018), Wragg et al. (2022), and Yadró Garcia et al. (n.d.). The sequencing reads (FASTQ files) were mapped to the reference honey bee genome Amel_HAv3.1 (Wallberg et al., 2019) using the Burrows-Wheeler Aligner (Vasimuddin et al., 2019). Variant calling and quality filtering were performed using the pipeline developed by Wragg et al. (2022), with further details provided in the supplementary information.

From the final VCF file, individuals with more than 30% missing genotypes were excluded. Similarly, SNPs with over 30% missing data or a minor allele frequency (MAF) below 0.5% were filtered out using PLINK 1.9 (Purcell et al., 2007).

We focused on the 46 CYP genes previously described by Claudianos et al. (2006) and recently curated by Dermauw et al. (2020), and 18 housekeeping genes (Lü et al., 2018; Wieczorek et al., 2020) within our haploid dataset (Table S2). We extended the target genes to include 1 kb upstream from the transcription start site to capture possible promoter elements (Haberle and Lenhard, 2016). SNPs retrieved from these regions were annotated using VEP v115.0 (McLaren et al., 2016) with the *A. mellifera* Amel_HAv3.1 as the reference genome, employing Sequence Ontology (SO) terminology to describe annotation features and properties (Eilbeck et al., 2005). To ensure consistency and avoid inflated variant counts due to multiple transcript isoforms, only Ensembl (Dyer et al., 2025) canonical transcripts were considered in the annotation process (Table S2).

### 2.3 Genetic diversity and population structure analyses

The haplotypes were created using a multi-step pipeline to process VCF files for specific populations and genes. Initially, VCF files were subsetted by population using a sample list from a population file, filtered for homozygous variants (1/1 genotypes) using bcftools, and converted to text and CSV formats to extract variant annotations. These were further filtered for missense SNPs and specific CYP genes using custom R scripts, which generate position-to-amino-acid change mappings. The filtered variants were used to create region-specific VCFs, from which haplotypes were constructed using the vcfR package in R by extracting genotypes, resolving them into haplotypes (0/1 for reference/alternate alleles), and counting unique haplotype sequences. Finally, haplotype maps were updated with amino acid change annotations, cleaned to remove missing data, and aggregated to retain the top two most frequent haplotypes per transcript across populations, producing output files used to generate images.

To evaluate patterns of genetic variation within and among subspecies and countries (including only those with ≥10 individuals), we estimated nucleotide diversity (π) (Nei and Li, 1979), Tajima’s D (Tajima, 1989), haplotype diversity (Hd) (Harris and DeGiorgio, 2017; Nei and Roychoudhury, 1974; Nei and Tajima, 1981) using in-house scripts implemented in R (4.4.1; R Core Team, 2014), and pairwise fixation index (F_ST_) for each subspecies and country using PLINK 1.9 (Purcell et al., 2007). For subspecies-level analyses, admixture proportions (Q-values) for each of the 1,467 individuals were previously inferred from genome-wide SNPs using ADMIXTURE, and only individuals with Q-values > 0.90 were retained (Henriques et al. 2018; Yadro-Garcia et al., 2026; Yadro-Garcia et al., unpublished data). In total, 428 individuals met this criterion, with sample sizes ranging from 70 in subspecies ligustica to 8 in subspecies caucasia (Table S11).

### 2.4 McDonald-Kreitman (MK) test

To investigate whether CYP genes have evolved under positive selection, we applied the imputed McDonald-Kreitman test as described by Rivera-Colón et al. (2026). Orthologous CYP coding sequences were identified across A. mellifera and A. cerana using OrthoFinder v3.0 (Emms and Kelly, 2019), resulting in 45 orthogroups (genes) for analysis. The imputed MK test was applied using the default derived allele frequency cutoff of 15% (Murga-Moreno et al., 2022). This test corrects for the confounding effect of slightly deleterious nsSNPs by estimating their contribution from the ratio of low-to-high frequency synonymous variants, using a derived allele frequency cutoff of 15%. Significance (per-gene) was assessed using Fisher’s exact test, with p-values adjusted for multiple testing using the Benjamini-Hochberg false discovery rate (FDR) procedure. Genes with an FDR < 0.05 were considered significant. Estimates of the proportion of adaptive substitutions (α) were obtained by pooling data across all 45 orthogroups.

### 2.5 In silico analysis of non-synonymous SNPs (nsSNPs)

The predicted (Alphafold) 3D structures of native proteins were retrieved from Uniprot (Bateman et al., 2025). The secondary structure elements, CYP9Q3 and CYP36A1, were annotated using SecStrAnnotator (https://sestra.ncbr.muni.cz) (Midlik et al., 2021) and visualised using PyMOL (Schrödinger, LLC, 2015).

To assess the functional impact of missense mutations, multiple predictive tools were applied, and their outputs were integrated into a scoring matrix to identify high-impact variants. In our scoring matrix, a score of 1 was assigned to any tool that classified a nsSNP as impactful, and 0 otherwise. For I-Mutant2.0 and DynaMut2 only extreme values (< -1.00 and > 1.00) were given a score of 1 (Frenz et al., 2020). Furthermore, because both tools predict changes in protein stability, their scores were averaged. The final score ranges from 0 (all tools predict neutral or no impact) to 5 (all tools predict high impact).

SIFT (Sorting Intolerant From Tolerant) served as the primary tool for nsSNP characterisation, predicting amino acid substitution effects based on sequence homology (Sim et al., 2012). SNPs with scores ≤ 0.05 were classified as affecting protein function, whereas those >0.05 were considered tolerated (Sim et al., 2012). To evaluate nsSNP effects on protein stability, we employed I-Mutant2.0 (Capriotti et al., 2005) (pH 7.0 and temperature 27°C) and DynaMut 2.0 (Rodrigues et al., 2020), which estimate free energy changes (ΔΔG) between wild-type and mutant proteins. ΔΔG < 0 indicates decreased stability, while ΔΔG > 0 indicates increased stability. BLOSUM62 (Henikoff and Henikoff, 1992) was employed to predict evolutionarily conserved amino acid residues, where positive scores indicate conservative substitutions and negative scores indicate non-conservative substitutions. Substitutions at highly conserved sites are more likely to impact protein function (Eddy, 2004; Henikoff and Henikoff, 1992). Missense-3D (https://missense3d.bc.ic.ac.uk/∼missense3d2/) (Ittisoponpisan et al., 2019) was used to flag potentially structurally damaging variants by examining multiple structural features (17 total). As this tool requires three-dimensional structural information, variants were analysed using AlphaFold-predicted models. DoGSite3, via ProteinsPlus, was employed for binding pockets prediction (Ehrt et al., 2025; Volkamer et al., 2012). Only the predicted pocket containing the Fe-binding cysteine residue was considered. Variants within these pockets may be particularly relevant, as changes in binding affinity can impact substrate affinity and, consequently, the metabolic capacity of CYP enzymes (Gyulkhandanyan et al., 2020; Hayward et al., 2023).

## 3. Results

### 3.1 Polymorphisms patterns in CYP and housekeeping genes

Screening of the 46 CYP genes, spanning over 0.29 Mb, yielded 5,756 SNPs, detected across the 1,467 individuals. In contrast, the 18 housekeeping genes, covering over 0.09 Mb, contained 1,728 SNPs. The SNP density was significantly higher in CYP genes, averaging one variant every 65.7 bp (± 50.45), compared with one every 106.5 bp (± 100.55) in housekeeping genes (p = 0.039) (Figures 2A and B, Table S3, File S1). The two gene groups exhibited similar transition-to-transversion (ti/tv) ratios: 5.54 for CYP genes and 5.38 for housekeeping genes. C/T transitions and T/A transversions were the most prevalent substitution types. As expected, coding sequence (CDS) regions displayed higher ti/tv ratios than intronic regions in both gene groups (CYP: 6.84 vs 5.12; housekeeping: 8.33 vs 4.88; Table S4).

**Figure 2.**
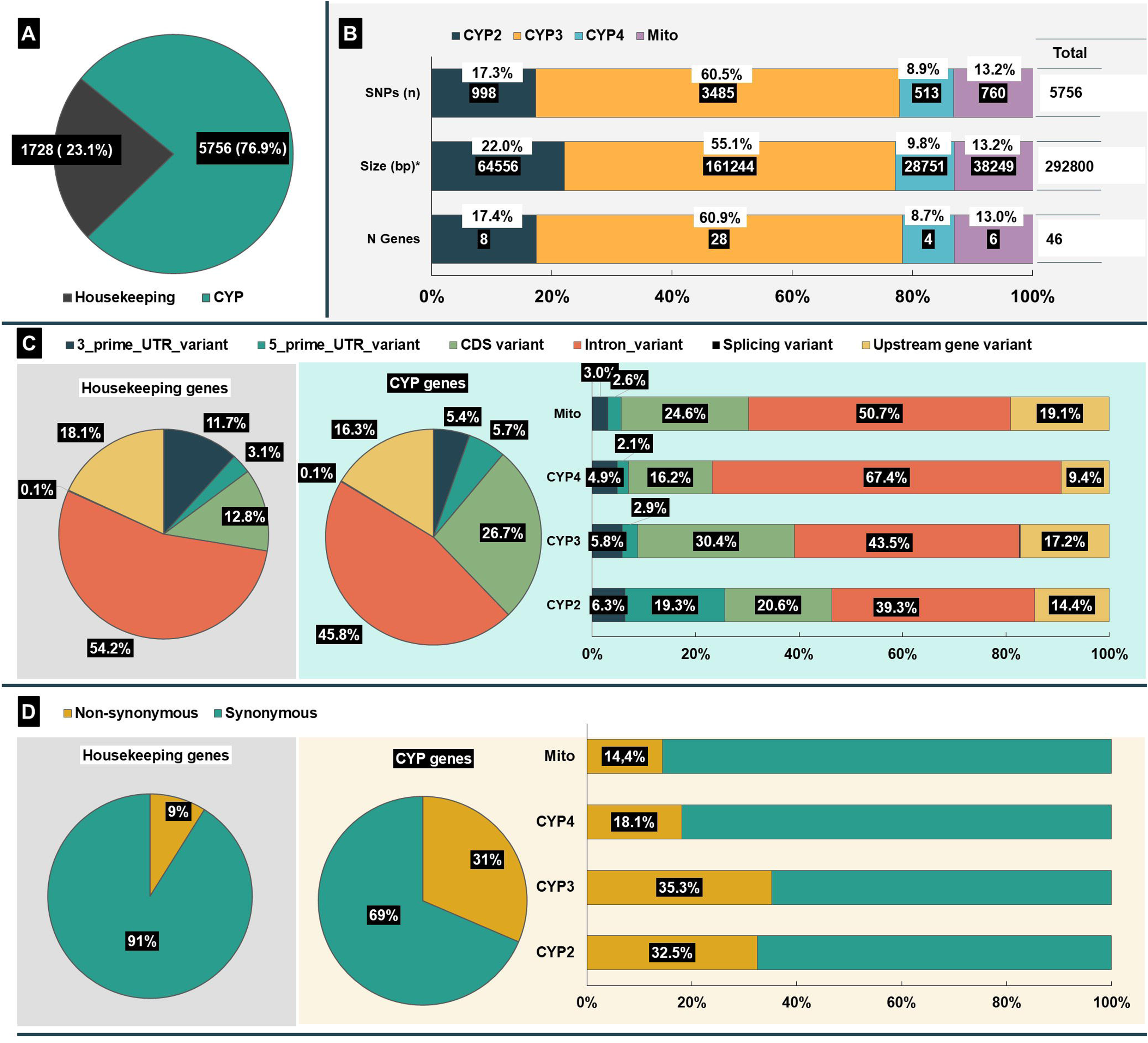
SNP analysis of CYP and housekeeping genes. A – Targeted screening identified 5,756 SNPs within the 46 CYP genes and 1,728 SNPs across the 18 housekeeping genes**. B –** SNPs breakdown by CYP clans. CYP3 clan contains the most variants (60.5%) and genes (60.9%). **C –** Variant type distribution among CYP genes, CYP clans, and housekeeping genes. CDS variant frequency is higher in CYP genes (26.7%) with the highest proportion in CYP3 (30.4%). **D –** Coding sequence variant distribution. Abbreviations: CDS – Coding Sequence variants. Mito – Mitochondrial CYP clan. * Size corresponds to the target region in our study: begins 1 kb upstream of the transcription start site of the canonical transcript and extends to its 3’ end.

SNPs distribution differed between the two gene groups. In CYP genes, intronic SNPs accounted for 45.8% of all SNPs, followed by coding SNPs (26.7%), of which 31.4% were non-synonymous. Housekeeping genes showed a higher proportion of intronic SNPs (54.2%), whereas coding SNPs comprised only 12.8%, with a predominance of synonymous substitutions (91.1%; Figures 2C and D).

The frequency of nsSNPs differed between the two gene groups: 483 and 20 for CYP and housekeeping genes, respectively (Figure 2D, Table S5). CYP genes accumulated significantly more nsSNPs per amino acid and more nsSNPs per synonymous SNPs than housekeeping genes (Mann-Whitney U, p < 0.001, File S2). The non-synonymous-to-synonymous (nsyn/syn) ratio in CYP genes ranged from 0.02 to 2.6, whereas in housekeeping genes, the ratio remained consistently low (0.0 – 0.009). Similarly, the non-synonymous-to-amino acid ratio (nsyn/aa) averaged 0.02 in CYP genes (Table S3), reaching up to 0.066 in some genes, but was below 0.009 in housekeeping genes (Table S5).

#### 3.1.1 CYP clans differ in SNP density and distribution

To investigate potential selective pressures and functional divergence among the four CYP clans, we examined SNP density and distribution patterns across CYP2, CYP3, CYP4, and mitochondrial genes. Given the functional heterogeneity of these clans, such patterns may offer further insights into their respective roles.

The distribution of CYP genes across clans was uneven. CYP3 constitutes the largest clan, comprising 28 genes, whereas CYP4 is the smallest, with only four genes (Dermauw et al., 2020; Feyereisen, 2006) (Figure 2B). SNP density varied among clans, although not significantly (File S1). CYP4 exhibited the lowest density, averaging one SNP per 102.6 bp (± 99.12), while CYP3 and mitochondrial clan genes showed higher densities, with one SNP per 60.7 (± 42.58) bp and 57.2 (± 39.16) bp, respectively (Table S3).

Ti/tv ratios in CDS also differed between clans, ranging from 6.11 in CYP3 to 12.8 in CYP4. In contrast, intronic regions displayed more moderate variation, with ti/tv ratios ranging from 4.76 in CYP2 to 5.78 in CYP4 (Table S4).

CDS variants were most prevalent in the CYP3 clan, accounting for 30.4% of its total variants. In contrast, only 16.2% of variants in CYP4 were located within coding regions (Figure 2C). The proportion of nsSNPs was highest in CYP3 and CYP2, at 35.3% and 32.5%, respectively, while the remaining clans did not exceed 20% (Figure 2D, Table S5).

The nsyn/syn ratio further distinguished the clans into two groups: CYP2 and CYP3 exhibited elevated ratios (mean of 0.76 ± 0.88 and 0.71 ± 0.61, respectively), whereas CYP4 and mitochondrial genes showed consistently lower ratios (0.30 ± 0.25 and 0.24 ± 0.19). This pattern was mirrored in the nsyn/aa ratio, with the highest mean in CYP3 (0.025 ± 0.02), and substantially lower in CYP4 (0.007 ± 0.005) and mitochondrial genes (0.008 ± 0.005) (Table S3). However, while the Kruskal-Wallis test detected significant differences among clans for nsyn/aa (p = 0.026), no significant differences were found for nsyn/syn (p = 0.050).

#### 3.1.2 CYP genes range from highly conserved to highly variable

Although CYP genes are classified into clans based on amino acid sequence similarity, reflecting shared structural and functional characteristics (Feyereisen, 2012), this phylogenetic grouping does not imply uniform selective pressures across clan members. Variant rate analysis revealed substantial heterogeneity, with outliers observed in CYP2 and mitochondrial genes (CYP303A1 and CYP302A1, respectively; Figure 3).

**Figure 3.**
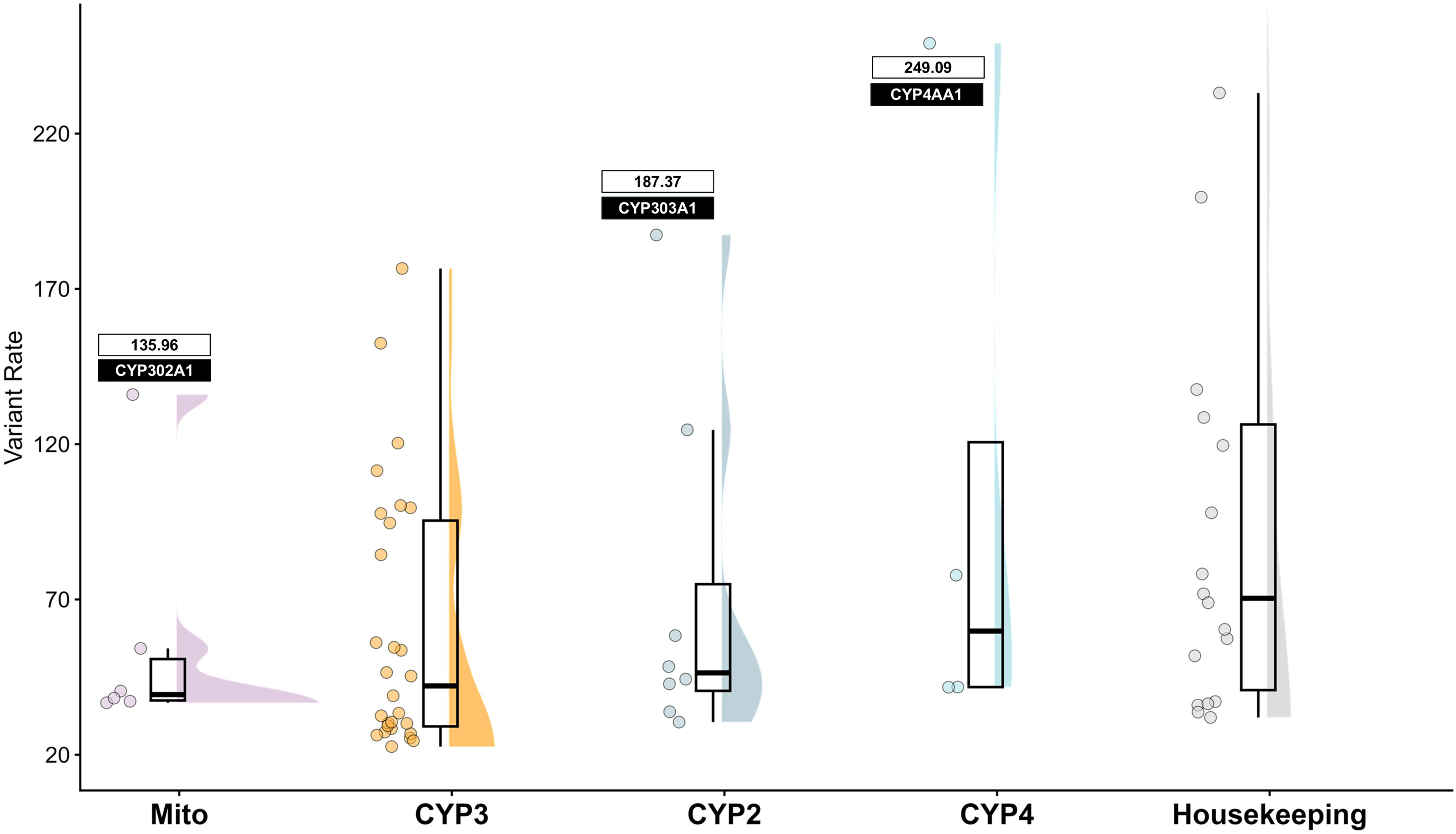
Distribution of variant rates across CYP clans in honey bees. Raincloud plots display the distribution of variant rates for each CYP clan, combining half-violin density plots (right), boxplots (center), and individual data points (left). Density curves are estimated via kernel density smoothing using the ggdist package in R and reflect the spread of the data within each clan. Boxplots indicate the interquartile range (IQR) with the median shown as a horizontal line. Individual genes are represented as jittered points, with fill colors corresponding to their respective clans. The y-axis is limited to 20–250 to focus on the main distribution; one outlier exceed this range: TATA-box-binding protein (Housekeeping, 436.7). Clans on the x-axis are ordered by median variant rate from lowest to highest. Variant rate is defined as the average distance (in bp) between variants. A variant rate of 50 indicates that, on average, one SNP is found every 50 base pairs. Lower values indicate higher variant density. Median variant rates are as follows: Mito (39.4), CYP3 (42.2), CYP2 (46.4), CYP4 (59.8), and Housekeeping (70.4).

To distinguish conserved from highly variable CYP genes, we examined two ratios: nsyn/syn and nsyn/aa (Figure 4). Housekeeping genes clustered near the origin, indicating low values for both ratios and suggesting strong purifying selection. Similarly, most CYP2, CYP4, and mitochondrial genes exhibited constrained variation, consistent with functional conservation (Figures 4A, C, and D). In contrast, several CYP2 and CYP3 genes exhibited elevated nsyn/syn ratios (> 1.0), including CYP6AS162, CYP6AS5, CYP369A1, CYP6AS1, CYP305D1, and CYP343A1 (Figures 4A and B).

**Figure 4.**
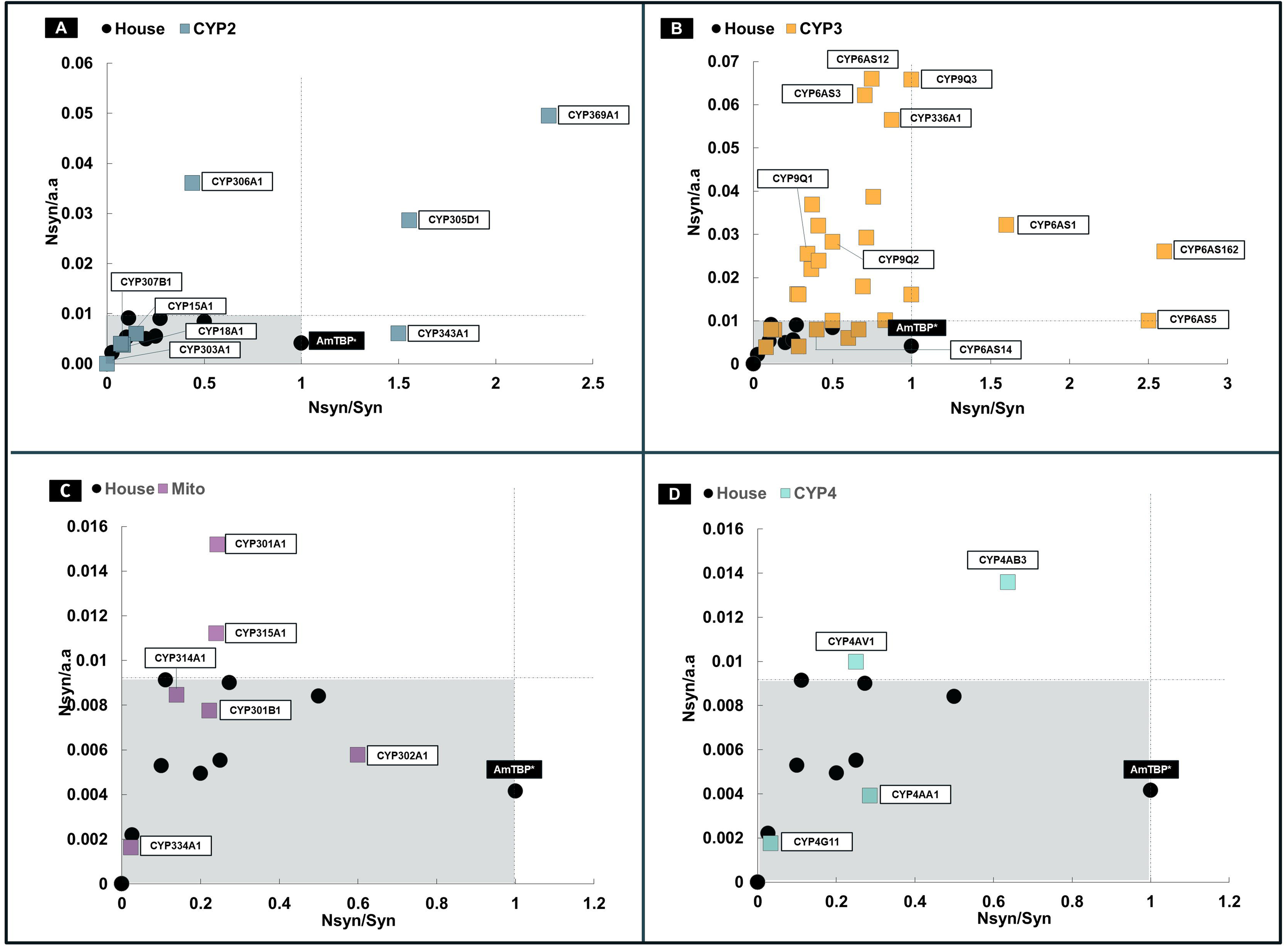
Non-synonymous mutational patterns across genes. **A –** CYP2 clan genes. **B –** CYP3 clan genes. **C –** Mitochondrial clan genes. **D –** CYP4 clan genes. Each square represents an individual CYP gene within the indicated clan, annotated with the corresponding encoded CYP enzyme. Housekeeping genes are clustered in the lower-left quadrant (grey area), characterized by low non-synonymous-to-synonymous (nsyn/syn) ratios and low non-synonymous mutations per amino acid (nsyn/a.a). Dashed lines indicate the upper thresholds for housekeeping genes for both nsyn/a.a and nsyn/syn ratios. In panel **B** (CYP3 clan), only selected genes are highlighted for clarity; complete information is provided in Table S5. AmTBP denotes the TATA-box-binding protein gene. Abbreviations: House, housekeeping genes; Mito, mitochondrial CYP clan. The asterisk marks the upper nsyn/syn ratio observed for housekeeping genes; however, this gene harbors only two coding sequence variants (one missense and one synonymous).

Notably, CYP3 genes (CYP6AS12, CYP9Q3, CYP6AS3, and CYP336A1) predominantly occupy the upper-left quadrant of the plot (characterised by high nsyn/aa but moderate nsyn/syn ratios), suggesting a distinct pattern of selective pressure (Figure 4B).

### 3.2 CYP Hd across genes, countries and subspecies

Haplotype frequency analysis can detect selective pressure, particularly when rare haplotypes occur at unusually high frequencies, a pattern that contrasts with the uniform frequencies expected for housekeeping genes. CYP genes, however, are known to accumulate multiple mutations that can influence enzymatic activity and substrate specificity (Zanger and Schwab, 2013). These linked polymorphisms are often indicative of adaptive responses to environmental pressures.

#### 3.2.1 Conservation and variability in CYP genes revealed by Hd

A total of 1,302 distinct haplotypes were identified, with a majority (84%, n = 1,094) located within the CYP3 clan (Table S6). This clan uniquely encompassed both the gene with the highest haplotype diversity (CYP9S1; Hd = 0.976) and the third least diverse (CYP6AS8; Hd = 0.068). In contrast, the CYP4 clan exhibited the lowest mean haplotype diversity across its members (Hd = 0.263 ± 0.296) (Tables S3 and S2) (Figure S2). Genes with higher diversity (Hd ≥ 0.9) are CYP9S1/CYP9R1, CYP6AS3, CYP6AS12, and CYP9Q3, all from the CYP3 clan (Table S5), which contain a higher nysn/aa (Figure 4B).

A comparison of the most frequent haplotype per CYP gene with the reference haplotype is shown in Figure 5. As expected, the reference haplotype was the most prevalent in most genes (27 out of 46). However, notable exceptions emerged: CYP6AS18 lacked any haplotype matching the reference, CYP6AS12 had only a single individual with reference haplotypes, and both CYP9S1/CYP9R1 and CYP6AS3 showed markedly low frequencies of reference haplotypes (n=38).

**Figure 5.**
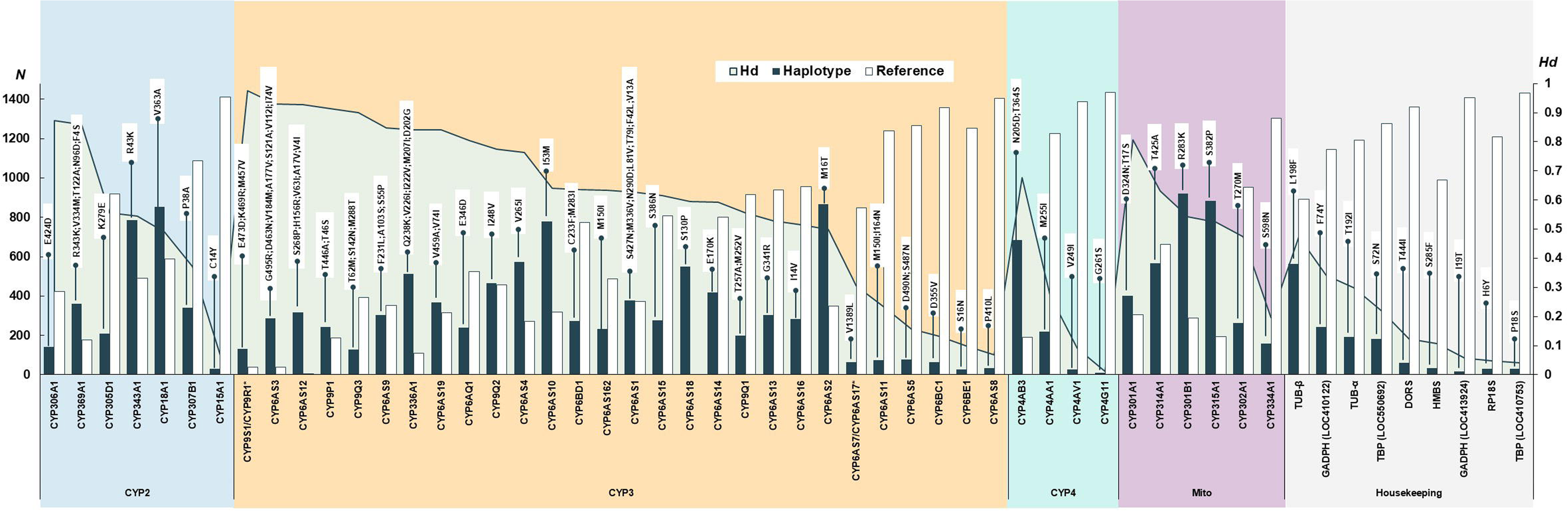
Haplotype frequency, distribution, and diversity across CYP clans and housekeeping genes. For each gene, only the most frequent missense haplotype (blue) and the reference haplotype (white) are shown. Haplotype diversity was lowest among CYP4 and housekeeping genes. Asterisks denote genes that, according to Dermauw et al. (2020), correspond to the fusion of two adjacent CYP genes. Abbreviations: Hd, haplotype diversity; TUB-β, tubulin beta chain gene; TUB-α, tubulin alpha chain gene; TBP, TATA-box-binding protein gene; GAPDH, glyceraldehyde-3-phosphate dehydrogenase gene; DORS, dorsal gene; HMBS, porphobilinogen deaminase gene; RP18S, 40S ribosomal protein S18 gene.

Reference haplotypes predominated among housekeeping genes (Figure 5). The mean haplotype diversity was low (0.19 ± 0.15; Table S5), and across nine genes with nsSNPs, 20 missense variants resulted in only 31 haplotypes. Most housekeeping genes harboured only two to four distinct haplotypes, underscoring their evolutionary constraints (Table S6). CYP genes exhibited significantly higher haplotype diversity than housekeeping genes (Mann-Whitney U, p < 0.001, File S3); however, no significant differences in Hd among CYP clan members were found (Kruskal-Wallis, p = 0.190).

#### 3.2.2 African and Middle Eastern countries show elevated CYP Hd

We analysed haplotype diversity and frequency by country (Table S7-10) and pure subspecies (Table S12-15, and Figures S3-5 illustrate the distribution of missing data across the analysed genes). Detailed results are presented in Tables S7-S10, showing haplotypes per gene and country. Significant differences were found for both groups (Kruskal-Wallis, p < 0.001, File S4).

Geographic haplotype diversity analysis revealed regional patterns across 19 countries (six countries were eliminated from the analysis as n < 10) (Figure 6A). Samples from Portugal, Spain, and Oman displayed the lowest median diversity for CYP genes (0.08 [0.0–0.34], 0.15 [0.0–0.466], and 0.16 [0.15–0.42], respectively), although the latter showed a notable exception: CYP9Q3 and CYP9S1/CYP9R1 (Hd = 0.85). In contrast, African and Middle Eastern populations – Morocco, Egypt, Jordan, Lebanon, and the UAE – along with Malta and Slovenia, consistently showed elevated diversity (Hd > 0.5). This geographic pattern persisted across most CYP clans (Figure S6). For CYP2 and CYP3, Portugal, Spain, Greece, and Oman clustered at the lower end of the Hd range, whereas African and Middle Eastern countries (Lebanon, Morocco, and the UAE for CYP2 and Egypt, Jordan, and the UAE for CYP3) dominated the upper end. CYP4 presented a distinct pattern: five countries (Cuba, Cyprus, Egypt, Lebanon, and Switzerland) exhibited a median Hd of 0 (with 0.327, 0.367, 0.726, 0.540, and 0.653 as the maximum values), whereas Slovenia (Hd = 0.30 [0–0.526]), Malta (Hd = 0.29 [0–0.506]), and the UAE (Hd = 0.28 [0 – 0.778]) displayed the highest values (Figure S6-C, File S4). Overall, the CYP2 and CYP3 clans exhibited the greatest haplotype diversity across all sampled populations.

**Figure 6.**
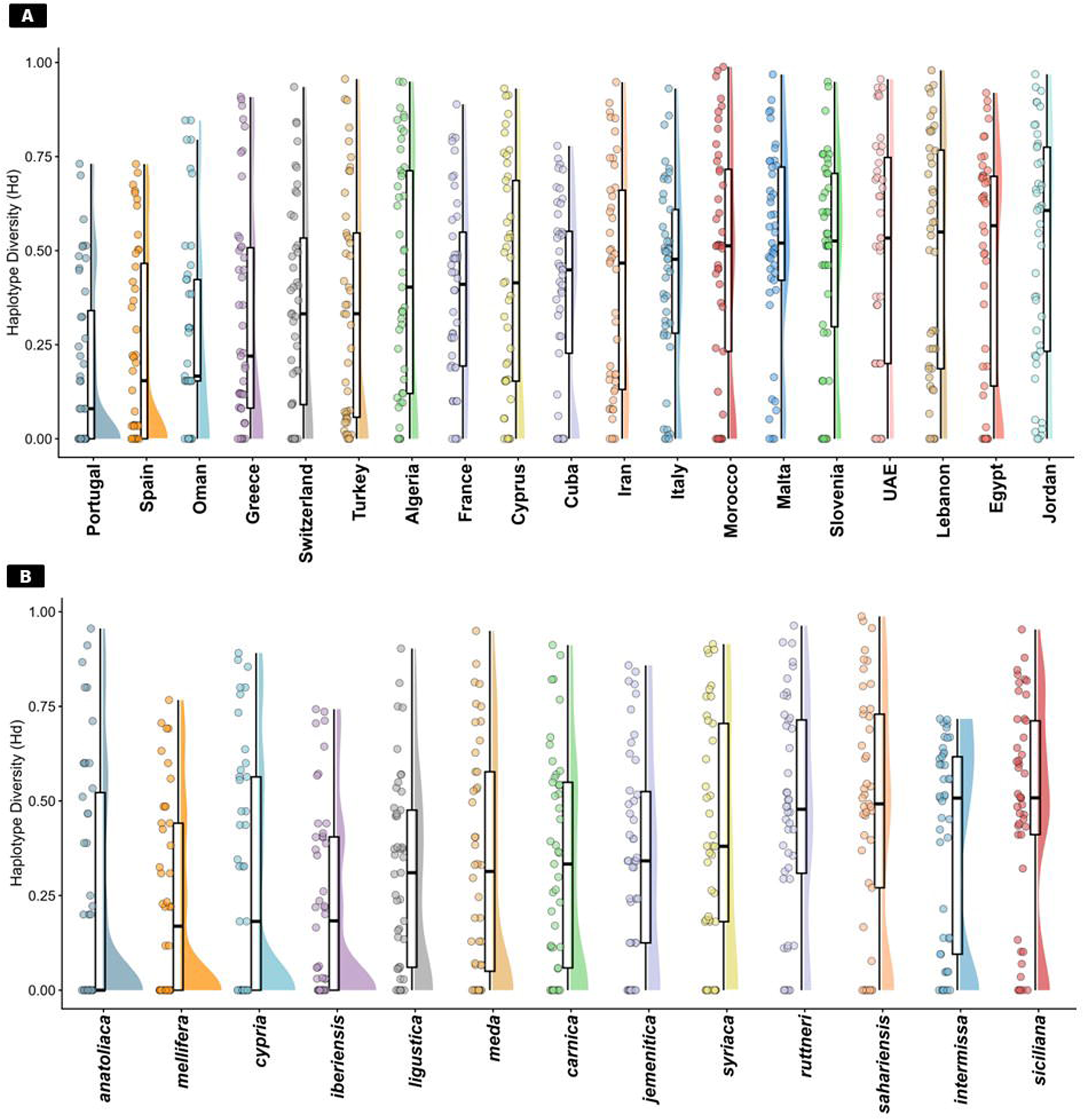
Haplotype Diversity (Hd). Haplotype diversity values are categorized by country **(A)** and pure subspecies **(B)**. Each country and subspecies’ distribution is visualized through three complementary elements: jittered data points (left) showing haplotype diversity for each CYP gene, boxplots (center, black outlines) displaying median, quartiles, and range, and half-violin density plots (right, colored) illustrating the probability distribution of Hd values. Countries and subspecies are ordered from lowest to highest median Hd (left to right), revealing substantial variation in genetic diversity. Pronounced differences in distribution shapes are evident, with Portugal and Spain (panel A), and A. m. anatoliaca, A. m. mellifera, A. m. cypria, and A. m. iberiensis (panel B) exhibiting strong skew, where most values concentrate at lower Hd. Furthermore, several countries and subspecies display extreme values approaching Hd = 1.0, primarily driven by elevated haplotype diversity in CYP3 clan genes. For detailed clan-specific patterns, see Figure S6, which decomposes these distributions by CYP clan.

#### 3.2.3 CYP Hd is greater in African *A. mellifera* subspecies

Our analysis identified 428 genetically pure individuals from 14 subspecies, with *A.m. ligustica* (16.4%), *A.m. iberiensis* (15.7%), and *A.m. siciliana* (13.3%) representing the largest groups (Table S11). Haplotype diversity analysis (Table S12-15, Figure 6B, File S4) revealed high mean CYPome values for *A. m. siciliana* (Hd = 0.51 [0.411 - 0.712]), *A. m. intermissa* (Hd=0.51 [0.095 – 0.617]) and *A. m. sahariensis* (Hd = 0.49 [0.271 – 0.729]). On the other hand, *A. m. mellifera* and *A. m. anatoliaca* displayed the lowest diversity (mean Hd = 0 [0 – 0.522] and Hd = 0.17 [0 – 0.441]). CYP clan analysis showed that CYP2 diversity peaked in *A. m. sahariensis* (mean Hd = 0.51 [0 – 0.871]), while it was low for *A. m. mellifera* (Hd = 0 [0 – 0.559] (Figure S7-A). Similarly, CYP3 reached maximum diversity in *A. m. siciliana* (Hd = 0.59 [0 – 0.953]), with *A. m. anatoliaca* ranking lowest (Hd = 0 [0 – 0.956] (Figure S7-B). CYP4 showed uniformly low diversity across all subspecies (Figure S7-C).

### 3.3. Nucleotide diversity

Nucleotide diversity (π) was assessed across CYP and housekeeping genes from 19 countries and 14 pure *Apis mellifera* subspecies (Figure 8). In both analyses, housekeeping genes displayed consistently low π values and showed no significant variation across groups (Kruskal-Wallis, country: p = 0.72, File S5; and subspecies p = 0.50,File S6), confirming their suitability as a baseline that reflects strong purifying selection, uniformly acting across countries and subspecies. CYP genes, in contrast, showed higher nucleotide diversity than housekeeping genes in all countries and subspecies. Statistical tests on CYP genes were highly significant (Kruskal-Wallis, p < 0.001), identifying significant pairwise differences among countries and subspecies (File S5 and S6).

**Figure 7.**
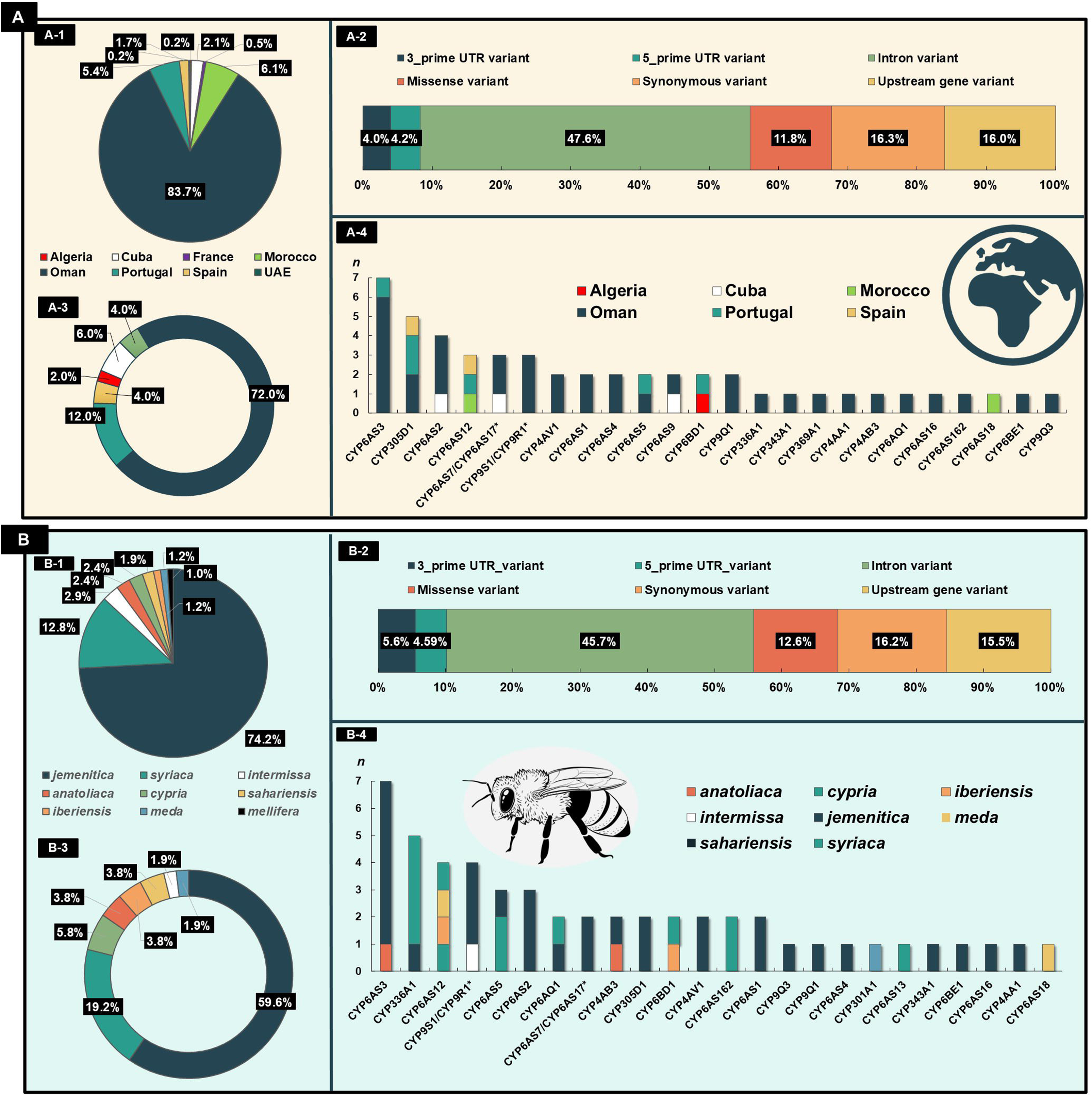
SNP analysis of CYP genes with FST over 0.9 by country and subspecies. **A –** Geographic-level differentiation. A-1: Proportion of countries containing variants with FST > 0.9. A-2: Distribution and classification of variant types. A-3: Distribution of missense variants by country. A-4: Distribution of missense variants across CYP genes and countries. **B –** Taxonomic-level differentiation. B-1: Proportion of subspecies containing variants with FST > 0.9. B-2: Distribution and classification of variant types. B-3: Distribution of missense variants by subspecies. B-4: Distribution of missense variants across CYP genes and subspecies. Bee illustration by Freepik (www.freepik.com).

**Figure 8.**
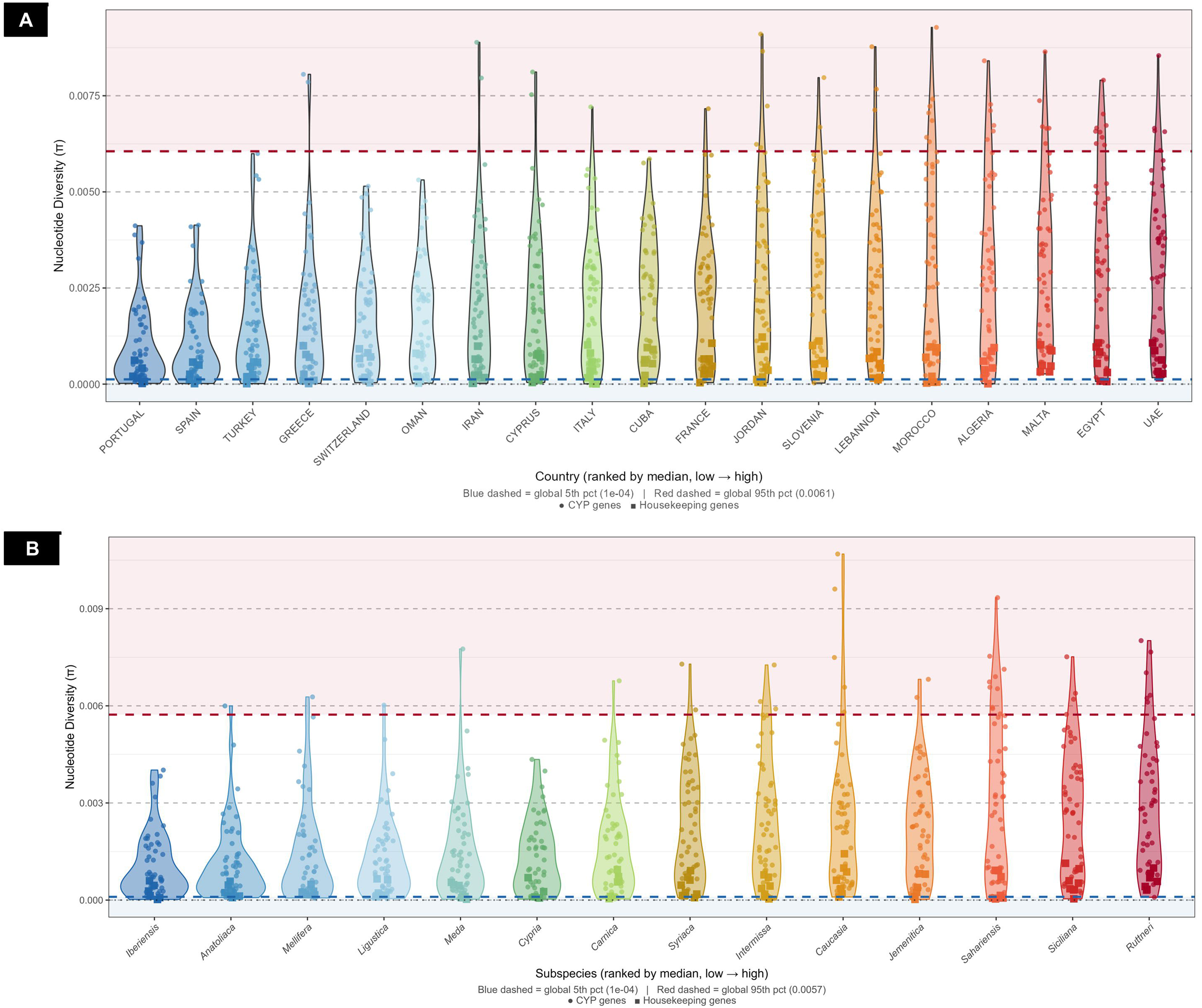
Nucleotide diversity for CYP and housekeeping genes genes across honeybee populations by country (A) and subspecies (B). Violin plots show the distribution of nucleotide diversity (π) for CYP genes (circles) and housekeeping genes (squares), per country (A) and subspecies (B). Median values are ranked from low to high. Blue and red dashed lines indicate the global 5th and 95th percentile thresholds, respectively (pink shading). Eastern Mediterranean and North African populations and subspecies generally display higher nucleotide diversity than their Western European counterparts.

Genes exceeding the global 95^th^ percentile across countries and subspecies were exclusively from the CYP3 clan (File S5 and S6), with one mitochondrial clan exception (CYP301A1 detected only in *sahariensis*). Ten genes surpassed the threshold in both analyses: CYP6AS16 was the most recurrent, appearing in 12 countries and eight subspecies, followed by CYP6AS1 (five countries, three subspecies), CYP6AS3 (five countries, two subspecies), and CYP9Q3 (three countries, four subspecies).

### 3.4 Tajima D’s selection signatures do not mirror nucleotide diversity

Housekeeping genes showed no significant variation across countries (Kruskal-Wallis, p = 0.84) (File S7) or subspecies (Kruskal-Wallis, p = 0.51) (File S8). And, once again there was significant differentiation across both countries and subspecies for CYP genes (p < 0.001).

Here, the geographic pattern of Tajima’s D did not mirror that of nucleotide diversity (Figure 9). Egypt, Malta, and Algeria showed the most positive median values [Q1-Q3] (0.172 [0.086 – 0.214], 0.154 [0.095 – 0.211], and 0.150 [0.079 – 0.199], respectively), whereas Oman, Turkey, and Switzerland were the most negative (medians − 0.085 [− 0.134 – 0.017], − 0.057 [− 0.118 – 0], and − 0.067 [− 0.176 – 0.092], respectively). At the subspecies level, *A.m. siciliana* and *A.m. ruttneri* had the most positive medians (0.149 [0.076 – 0.269] and 0.116 [0.069 – 0.181], respectively), whereas *A.m. ligustica* and *carnica* were the most negative (− 0.091 [− 0.203 – 0.102] and − 0.034 [− 0.134 – 0.068], respectively).

**Figure 9.**
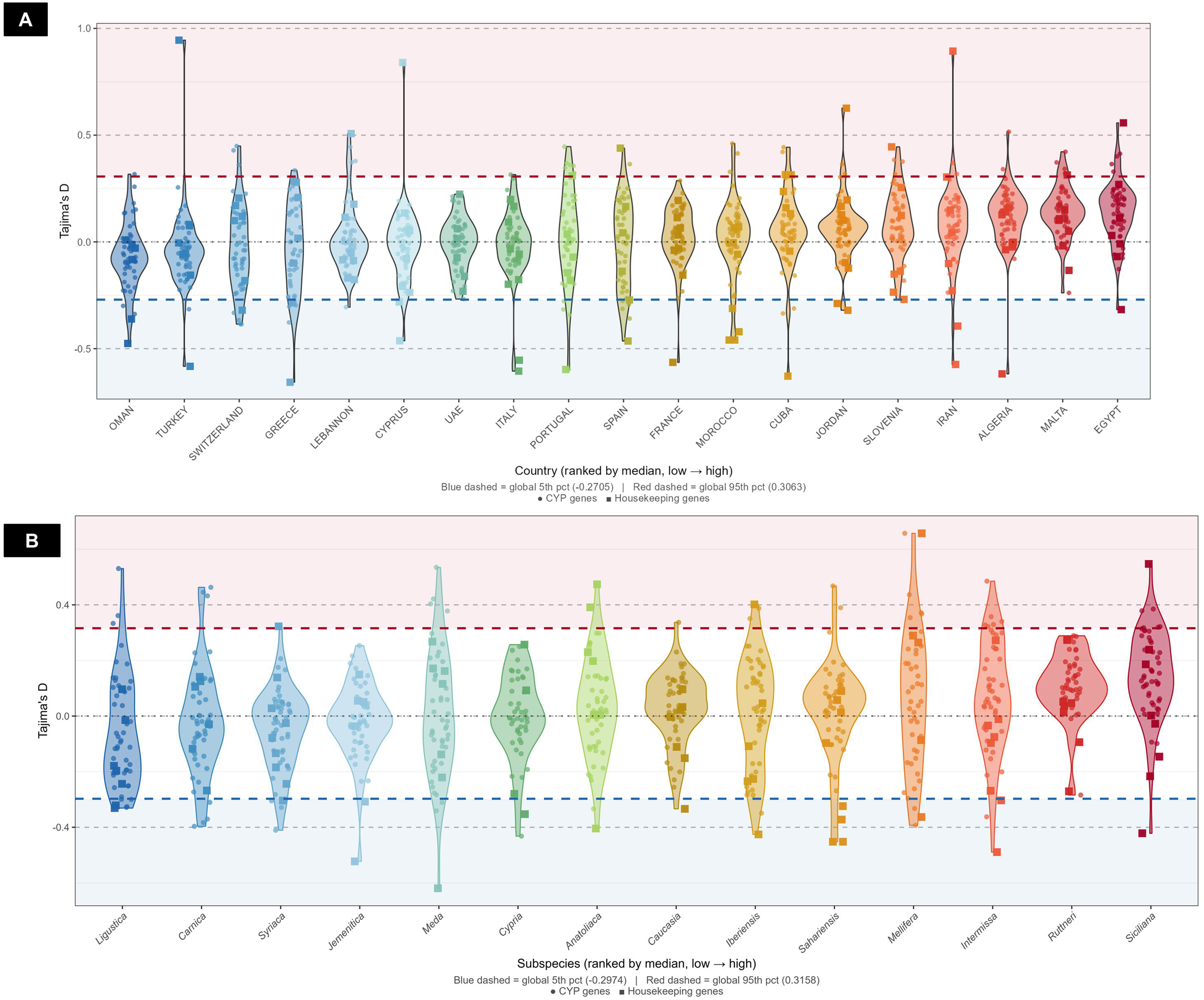
Tajima’s D of CYP and housekeeping genes across honeybee populations by country (A) and subspecies (B). Violin plots show the distribution of Tajima’s D for CYP genes (circles) and housekeeping genes (squares), ranked by median from low to high. Blue and red dashed lines indicate the global 5th and 95th percentile thresholds, respectively, with blue and pink shading highlighting values below and above these bounds. Most populations cluster near neutrality (D ≈ 0), though populations from Oman and Turkey show a negative skew suggesting purifying selection or population expansion, while Egypt, Malta, and Siciliana and Ruttneri subspecies tend toward positive values, indicative of balancing selection or demographic contraction.

Genes at the extreme tails (95^th^ and 5^th^ percentiles) were identified (File S7 and S8). CYP genes most consistently in the negative tail (both country and subspecies) included CYP6BD1, CYP6AS4, CYP6AS14, CYP303A1, CYP4AA1, and CYP301A, whereas the positive tail was dominated by CYP3 clan genes (CYP302A1, CYP6AS1, CYP6AS16, CYP343A1, and CYP9P1). CYP6BD1, CYP6AS4, CYP6AS14, and CYP4AB3 appeared in both tails depending on the subspecies or geographic context. For housekeeping genes*, LOC410753* (*TATA-box-binding protein*) was the most recurrent in the negative tail (eight countries, four subspecies).

### 3.5 CYP3 clan genes as hotspots of population differentiation

F_ST_ analysis revealed pronounced genetic differentiation. A total of 414 and 424 highly differentiated SNPs (F_ST_ > 0.9) were identified across 46 CYP genes in subspecies and country analyses, respectively. Most of the variants (n = 299, 72%) were shared between both datasets (File S9).

*A. m. jemenitica* and Oman emerged as the most genetically distinct populations, contributing with 74% and 84% of variants in the subspecies and country analyses, respectively. Given that the Omani samples consisted exclusively of *A. m. jemenitica,* this was expected. The remaining 107 SNPs were distributed across *A. m. syriaca (53)*, *A. m. intermissa (*12), *A. m. anatoliaca (10)*, *A. m. cypria (10)*, *A.m. sahariensis (8)*, *A. m. iberiensis (5)*, *A. m. meda (5)*, and *A. m. mellifera* (4). Seven additional countries (Algeria, Cuba, France, Morocco, Portugal, Spain, and the UAE) also harboured highly differentiated SNPs, although at substantially lower frequencies (Figure 7A, Table S16).

CYP3 clan genes were significantly enriched in both subspecies (OR = 1.53, P = 0.004) and country (OR = 1.68, P = 0.0003) analyses, accounting for 64.7% and 67.07% of the variants in each dataset, respectively. In contrast, CYP2 clan genes were significantly depleted (subspecies: OR = 0.60, p = 0.006; country: OR = 0.47, p = 0.0001) (File S9).

Remarkably, CYP6BE1 and CYP6AS3 carried the greatest number of highly differentiated SNPs in the subspecies (n = 25) and country (n = 38) analyses.

### 3.6 Evidence for adaptive selection

The imputed MK test revealed strong evidence of positive selection in *A. mellifera* using *A. cerana* as an outgroup (α = 0.559, p < 0.001), indicating that approximately 56% of nsSNPs in CYP genes were driven by positive selection (File S10). The strongest signal was detected in most CYP3 clan genes. In particular, for CYP6AS16 (α = 0.911, p_adj = 0.004) followed by CYP6AS11, CYP6AS7/CYP6AS17 (both α = 1.000), CYP6AS4 (α = 0.728) and CYP9Q2 (α = 0.822). DoS was positive for all FDR significant genes (Figure 10).

**Figure 10.**
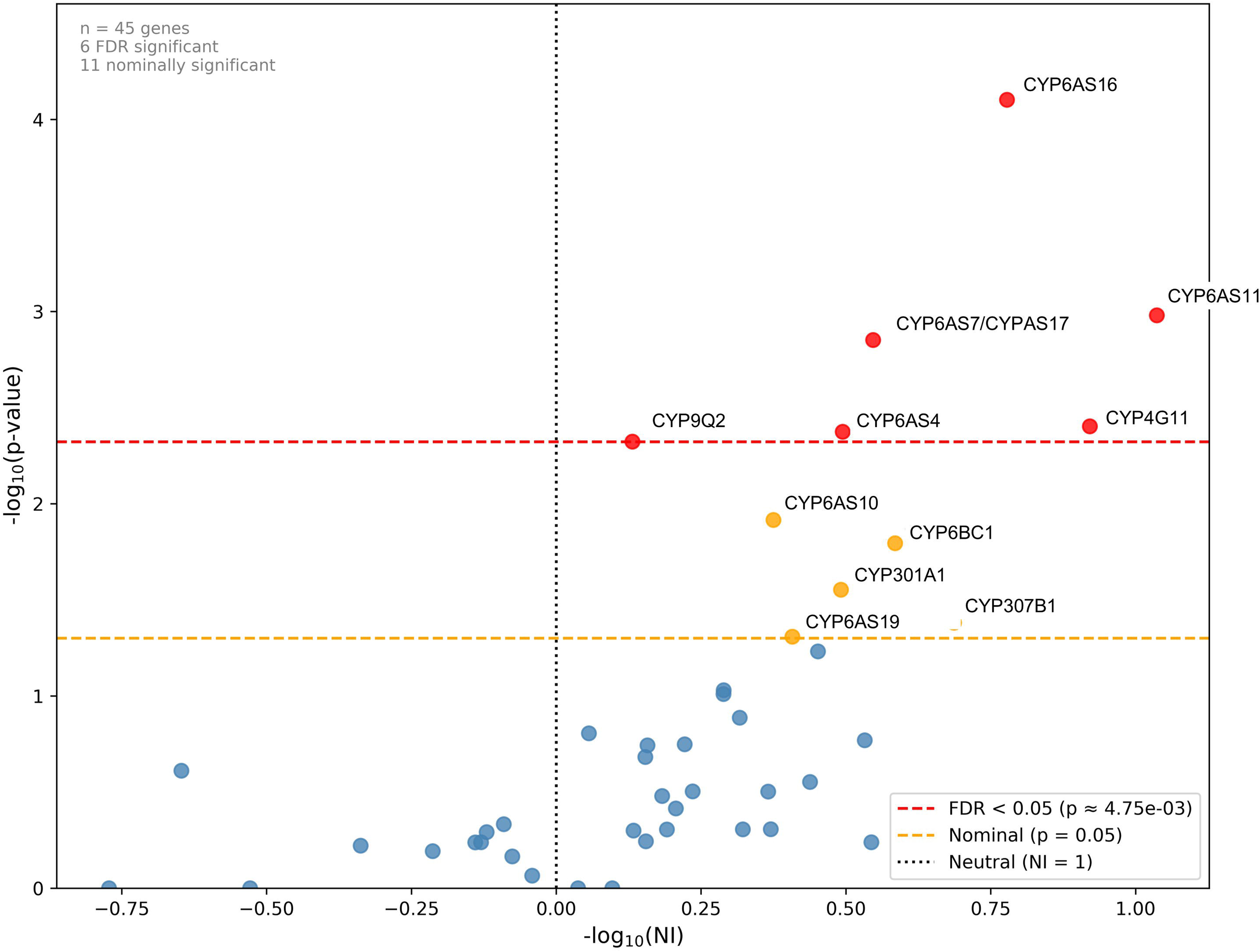
McDonald-Kreitman test results for 45 CYP genes in Apis mellifera versus Apis cerana. Volcano plot displaying the neutrality index (NI, log□□-transformed) against statistical significance (−log□□ p-value) for each gene. Points to the right of the vertical dotted line (NI < 1) indicate genes with an excess of nonsynonymous polymorphism relative to divergence, suggesting relaxed or negative selection, while points to the left (NI > 1) suggest positive selection. Red points exceed the FDR < 0.05 threshold (red dashed line; p = 4.75×10⁻³) and orange points are nominally significant (p < 0.05, orange dashed line). Of 45 genes tested, 6 were FDR-significant and 11 nominally significant, with CYP6AS16, CYP6AS11, CYP6AS7/CYPAS17, and CYP4G11 among the top candidates showing evidence of adaptive evolution.

### 3.7 *In silico* analysis of nsSNPs

The structural impact prediction of nsSNPs, conducted only on the 479 missense variants found for CYP genes, is summarised in Table S18. CYP3 and mito clan genes showed the highest mean impact scores (0.78 ± 0.79 and 0.83 ± 0.96, respectively) among the studied clans, while CYP4 genes displayed the lowest score (0.63 ± 0.52), indicating fewer structural alterations. High-impact SNPs (scores ≥ 3.5) were rare, occurring only three times. No variant reached the maximum possible score of 5. The highest observed impact score of 4 was recorded for the G302D SNP in CYP9S1/CYP9R1 and the R175G variant in CYP6AS162. Most SNPs were low impact with no statistical significance between clans or genes Figures 11 and 12 present the allele frequencies and estimated impact scores for CYP9Q3 and CYP336A1 SNPs, along with the annotations of relevant secondary structure elements. CYP9Q3 contains 33 missense variants, with nearly half (48.5%, n = 16) located in loop regions and a mean mutational score of 0.71 ± 0.59. The T62M variant stands out as having both the highest impact score (2.0) and the highest allele frequency (0.43). CYP336A1 displayed a comparable variant distribution, with 28 missense variants and a mean mutational score of 0.80 ± 0.81. Most variants (39.3%, n = 11) were again concentrated in loop regions. Although five variants exhibited relatively high allele frequencies (0.49 – 0.60), their impact scores remained low (0 – 1) (Figure 12).

**Figure 11.**
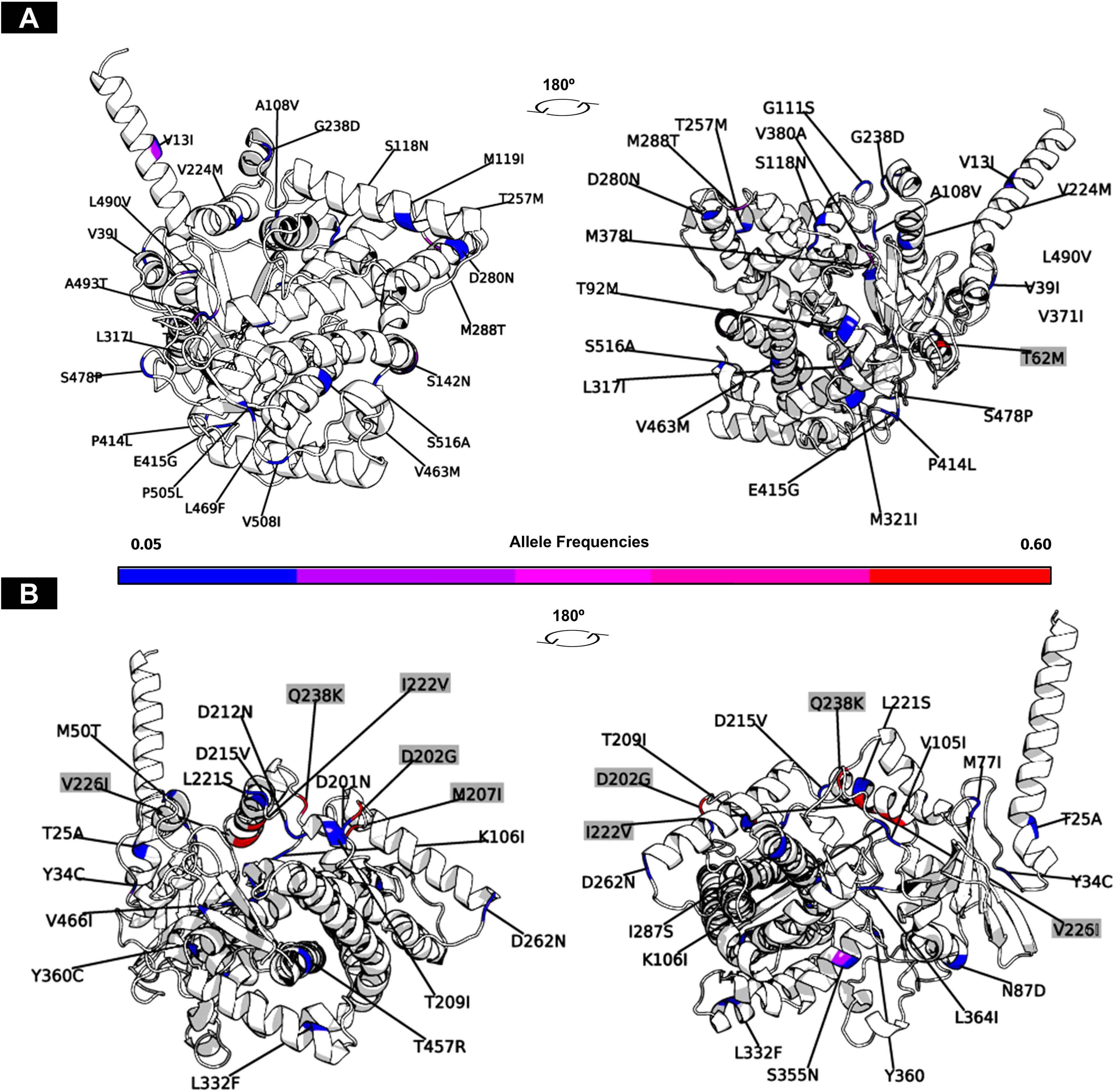
CYP9Q3 and CYP336A1 missense SNP distribution and allele frequency. A – CYP9Q3, B- CYP336A1. Variants with the highest allele frequency are highlighted: T62M (0.43), D202G (0.59), I222V (0.53), V226I, M207I (both 0.52) and Q238K (0.49).

**Figure 12.**
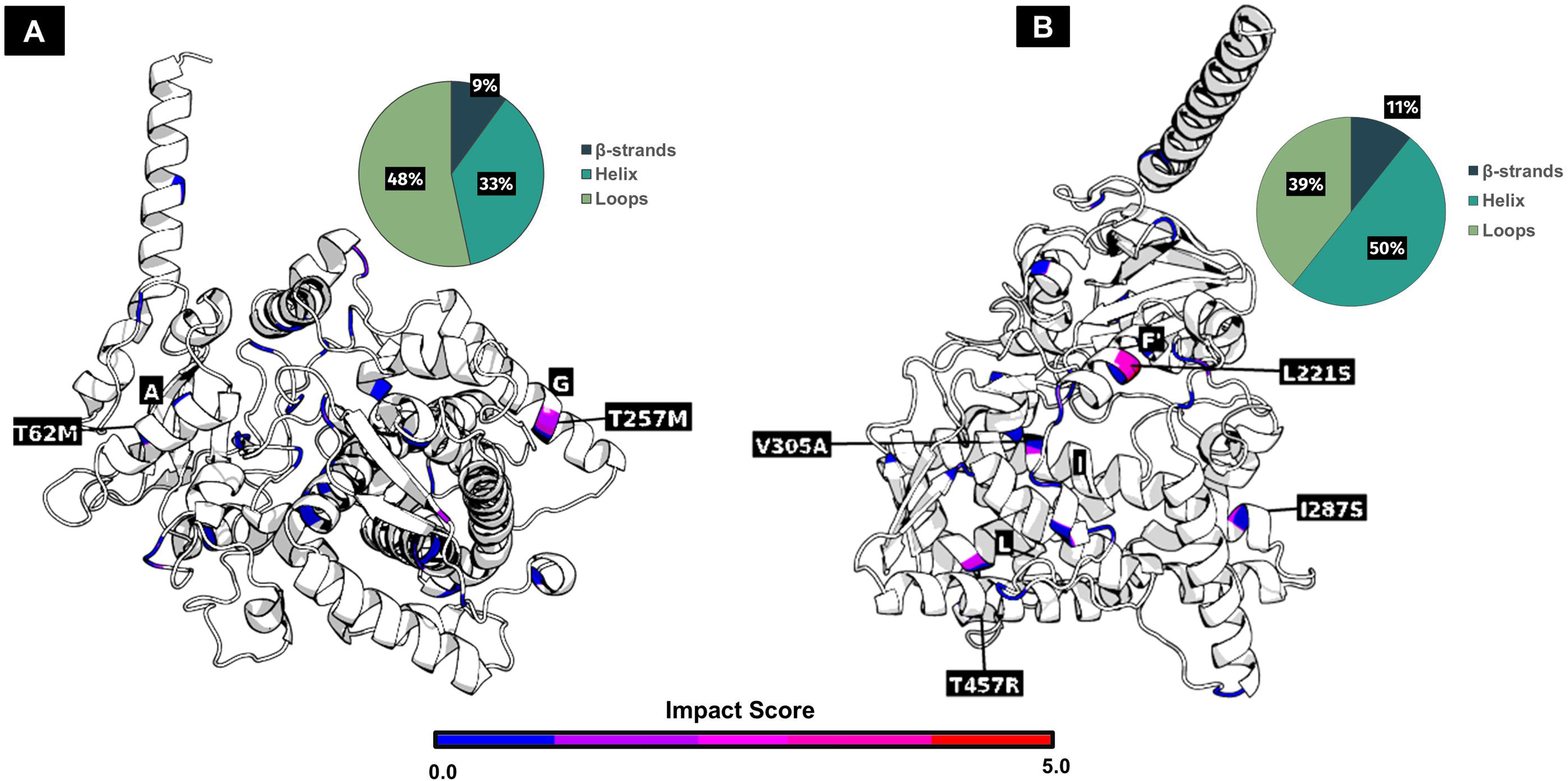
CYP9Q3 and CYP336A1 missense variants. A – Impact scores of variants and proportion of missense SNPs across secondary structures of CYP9Q3. B – Impact scores of variants and proportion of missense SNPs across secondary structures of CYP336A1. Variants with the highest impact scores (and their corresponding helices) are highlighted for each protein: T62M and T257M (both 2.0) in CYP9Q3; L221S (3.0), I287S, V305A and T457R (2.0) in CYP336A1.

## 4. Discussion

### 4.1 A foundational catalogue for honey bee pharmacogenomics

Cytochrome P450 monooxygenases (CYPs) are central to insect detoxification, enabling the metabolism of plant allelochemicals and pesticides (Feyereisen, 2012). Despite their importance, natural variation in honey bee CYP genes remains poorly characterised. Drawing a parallel with human medicine is instructive: modern pharmacogenetics emerged from the characterisation of CYP polymorphisms and their functional impacts (Zanger and Schwab, 2013).

Discoveries showing how variants in CYP2D6, CYP2C9, and CYP3A4 alter drug metabolism have revolutionised personalised medicine, allowing clinicians to anticipate drug efficacy and toxicity based on individual genotypes (Zanger and Schwab, 2013; Zhou et al., 2009).

This study provides the first comprehensive catalogue of CYP genetic variation across *A. mellifera*subspecies– comprising 5,756 SNPs identified in the 46 CYP genes of 18 subspecies sampled across 25 countries. A resource such as this dataset will enable researchers to leverage naturally occurring variation for *in silico* approaches, including molecular docking studies using platforms like the Insect cytochrome P450 Database (http://www.insectp450.net/) (Wu et al., 2025) and molecular dynamics simulations to predict how specific haplotypes alter substrate binding or catalytic efficiency.

The biological relevance of this dataset is validated by significant geographic and subspecies differentiation in haplotype diversity and is further cemented by strong evidence of positive selection acting on the CYP gene family (imputed McDonald–Kreitman test, α = 0.559, p < 0.001), in which approximately 56% of nsSNPs were driven by adaptive evolution.

Furthermore, the recent publication of an *in silico* pipeline for arthropod P450 model generation and active site docking analysis by Hayward et al. (2026) enables the immediate exploitation of our dataset by these computational tools.

In the spirit of human pharmacogenomics, this SNP catalogue contributes to the growing field of insect toxicogenomics, aiming to enhance the understanding of pesticide responses and improve the health and resilience of this vital pollinator.

### 4.2 Differential selective pressures shape CYP clan diversity

CYP genes in honey bees (and insects alike) bridge core physiological functions, such as the biosynthesis and metabolism of pheromones, cuticular hydrocarbons, and hormones, with the detoxification of foreign compounds. Consequently, few *A. mellifera* CYPs substrates are known. Furthermore, the functional characterization of specific CYP genes is complicated by the diversity of roles they assume (Feyereisen, 2012).

The higher proportion of non-synonymous variants (compared with other clans) in the CYP2 and CYP3 families, as shown in Figures 2D, 4A, and 4B, suggests that these genes are evolving under diversifying selection. This is consistent with their established roles in xenobiotic detoxification, where genetic variability may confer adaptive advantages in response to environmental pressures, such as exposure to plant-derived toxins and pesticides (Feyereisen, 2012). For instance, the CYP9Q subfamily within the CYP3 clan has been directly implicated in the detoxification of key classes of insecticides, including neonicotinoids, organophosphates, diamides, butenolides and pyrethroids, (Bass et al., 2024; Haas et al., 2022a, 2021; Manjon et al., 2018; Mao et al., 2011). The elevated non-synonymous variation observed in the CYP9Q genes may enhance their functional diversity, potentially increasing the adaptive potential of honey bee populations in agricultural landscapes where exposure to pesticides may be high.

In contrast, the CYP4 family and mitochondrial CYPs exhibited markedly different patterns, with a predominance of synonymous substitutions and lower non-synonymous variation (Figures 4C and D). This suggests that these genes are under purifying selection, likely because of their critical roles in essential physiological processes. CYP4 genes are primarily involved in cuticular hydrocarbon synthesis and fatty acid metabolism, which are vital for maintaining the structural integrity of the exoskeleton and energy homeostasis (Qiu et al., 2012). Similarly, mitochondrial CYPs contribute to core metabolic pathways, and even minor disruptions can have severe fitness consequences (Feyereisen, 2012; Rewitz et al., 2007). The reduced non-synonymous variation in these gene families reflects the evolutionary constraints imposed by their important physiological roles, which limit their tolerance to protein-altering mutations.

This clan-level contrast is also reflected in nucleotide and haplotype diversity patterns. CYP3 clan Hd and π differed significantly across both countries and subspecies. In contrast, CYP4 showed no significant differentiation. This is consistent with the hypothesis that CYP3 genes respond to local selection pressures, whereas CYP4 genes are constrained by core physiological processes that do not vary with the environment (Figure S2). The mitochondrial clan is more nuanced. It showed significant geographic and taxonomic differentiation in Hd and the highest DoS (0.135) and mean α (0.739) across all clans in the MK analysis, suggesting that positive selection has acted on mitochondrial CYPs despite their conserved core functions. However, it is worth remembering that no gene in this clan reached the FDR significance level. For CYP2, only 50% of the genes showed positive DoS values, and the mean α was negative (-0.153). Both highly conserved (CYP303A1, nsyn/syn = 0) and apparently rapidly evolving genes (CYP369A1, nsyn/syn = 2.27) suggest thatt this clan is not functionally uniform.

### 4.3 Gene-specific evolutionary signatures in the CYP3 clan

The high haplotype diversity within CYP3 (84% of all haplotypes and 1,094 distinct variants) is remarkable and reflects the central role of this clan in xenobiotic metabolism. Interestingly, many of the most variable and highly differentiated genes in the CYP3 clan belong to the CYP6 family, which is specifically associated with insecticide resistance in multiple insect orders (Han et al., 2022; Li et al., 2007b; Müller et al., 2007; Xiao and Lu, 2022; Zou et al., 2019). The CYP6 family has undergone extensive gene duplication and diversification in insects, creating a reservoir of catalytic diversity for metabolising structurally diverse compounds (Feyereisen, 2012).

Feyereisen (2012) noted that most CYP3 genes fit Berenbaum (2002) concept of “environmental response genes”: high diversity, frequent duplication, rapid evolutionary rates, genomic clustering, and tissue-or stage-specific expression. The CYP6AS subfamily provides a paradigmatic example. Unique to Hymenoptera, this subfamily underwent rapid evolutionary diversification and is the largest CYP subfamily in the *A. mellifera* genome, comprising 15 of the 46 CYPome genes (Claudianos et al., 2006). This lineage-specific “bloom” is thought to reflect selective pressures associated with secondary plant metabolite detoxification (Feyereisen, 2012). Flavonoids (particularly quercetin) found in both pollen and honey (Campos et al., 1997) are mainly metabolised by CYP6AS enzymes (CYP6AS1, CYP6AS3, CYP6AS4, and CYP6AS10) (Mao et al., 2009). However, our results, consistent with previous studies (Fujita et al., 2018; Wu et al., 2020), caution against broad generalisations.

Several CYP6AS genes exhibit patterns of genetic variation inconsistent with the expectations for enzymes involved in detoxifying dietary phytochemicals. This is particularly relevant given that honey is derived from diverse floral nectars, and its phytochemical profile varies across space and phenological cycles. We found that CYP6AS8, CYP6AS11, CYP6AS14, and CYP6AS16 exhibited polymorphism levels comparable to those of housekeeping genes. Specifically, these genes cluster in regions characterised by low nsyn/syn and low nsyn/aa (Figure 4B, Table S5). CYP6AS8 and CYP6AS11 also show very low haplotype diversity (Figure 5, Hd = 0.068 and 0.231, respectively). Moreover, CYP6AS8 is seemingly involved in fatty acid hydroxylation (Wu et al., 2020). The mutational patterns we observed for CYP6AS8 are consistent with an enzyme performing essential, conserved physiological functions rather than xenobiotic detoxification. Notably, these two genes are among the most highly expressed P450s in queen and worker mandibular glands, respectively (Wu et al., 2017), suggesting a role in mandibular gland secretion biosynthesis.

This illustrates how gene duplication can lead to functional divergence – while some CYP6AS genes have retained or evolved xenobiotic functions, others have become specialised for essential biosynthetic roles (Mao et al., 2009; Wu et al., 2020). CYP6AS11 is an interesting case. It was FDR-significant in the MK test, indicating a strong history of adaptive fixations between *A. mellifera and A. cerana*. This dissociation between low intraspecies polymorphism and high interspecies divergence points to positive selection having swept the gene to fixation within the mellifera species.

For CYP6AS14, gene expression data are contradictory: the gene has been reported to be upregulated following exposure to imidacloprid (Wu et al., 2017), thymol (Boncristiani et al., 2012), acrinathrin, stearine, and chlorpyrifos-ethyl (El Agrebi et al., 2024), but downregulated in response to thiamethoxam (Tesovnik et al., 2020), imidacloprid, and coumaphos (Chaimanee et al., 2016). Our data (showing low genetic diversity and constrained variation patterns) do not support a primary or direct role for CYP6AS14 in xenobiotic metabolism. To date, no experimental evidence supports the classification of this enzyme as a xenobiotic metaboliser. Furthermore, Tajima’s D results for this gene show no consistent directional signal having appeared in both positive and negative tails. This is a pattern more consistent with demographic heterogeneity than with strong purifying or positive selection.

These observations highlight an important caveat of expression-based studies: the induction of gene expression alone does not translate into direct involvement in detoxification. For CYP genes, because they participate in a wide range of essential physiological processes, pesticide exposure is expected to broadly alter their expression as part of a generalised stress response rather than a targeted detoxification mechanism. Additional support for this interpretation comes from reports that imidacloprid-exposed larvae exhibit disrupted lipid homeostasis (Derecka et al., 2013). Thus, the up- and downregulation observed in previously mentioned studies may reflect indirect effects of pesticide exposure on hormone and metabolism-related pathways, potentially contributing to reduced lifespan and sublethal effects rather than direct xenobiotic metabolism.

CYP6AS5 presents an interesting case: low nsyn/aa, high nsyn/syn ratio, and one of the lowest haplotype diversities within the CYP3 clan: consistent with very limited relevance in detoxification. Manjon et al. (2018) reported that CYP6AS5 is a weak thiacloprid metabolizer, suggesting a marginal role in xenobiotic detoxification. Intriguingly, Li et al. (2021) observed that larvae exposed to thiamethoxam showed increased juvenile hormone levels accompanied by upregulation of CYP6AS5, leading them to propose a potential role in hormonal regulation rather than xenobiotic detoxification. The MK test yielded a non-significant result (α = 1.00, p = 1.0), and CYP6AS5 showed the most negative DoS among all CYP3 genes (- 0.414). Here, the near absence of nonsynonyous polymorphism (relative to divergence) is artificially inflating α, suggesting that CYYP6AS5 may be under strong purifying selection.

### 4.4 CYP9Q subfamily and CYP336A1: specialized xenobiotic metabolizers

Enzymes of the CYP9Q subfamily are the xenobiotic metabolizers of excellence. Mao et al. (2011) demonstrated through heterologous expression that CYP9Q1, CYP9Q2, and CYP9Q3 can metabolize tau-fluvalinate, an acaricide widely used for *Varroa destructor* control. Expanding on this finding, Manjon et al. (2018) used a combination of functional assays and toxicity tests to show that CYP9Q enzymes, especially CYP9Q3, play a crucial role in detoxifying neonicotinoids. These findings support a model in which CYP9Q3 acts as the first line of defense in honey bees against both acaricides and neonicotinoids, highlighting its status as a key detoxifier.

Consistent with this function, one would expect the gene to maintain high genetic diversity to cope with the wide range of environmental toxins encountered by honey bee foragers. Indeed, the elevated Hd and nsyn/aa observed for CYP9Q3 aligns with the selection signature expected for a major detoxification enzyme. In contrast, CYP9Q1-2 showed more constrained genetic variation. CYP9Q2 was FDR significant in the MK test (α = 0.822, DoS = 0.069, p_adj = 0.036), whereas CYP9Q3 had a slightly negative DoS (-0.024) and non-significant α. This suggests complementary strategies within the subfamily: CYP9Q3’s extreme polymorphism may reflect diversifying selection, whereas CYP9Q2 has accumulated adaptive fixation between species.

CYP336A1 has recently been shown to metabolise nicotine with a key residue (D298 in helix I) functioning as a key determinant of metabolic action against nicotine and atropine, suggesting specialisation toward alkaloid detoxification (Haas et al., 2023). The observed enrichment of variants in the flexible structural elements (loops) of CYP336A1 and CYP9Q3 suggests that both enzymes tolerate amino acid substitutions that modulate substrate interactions without compromising the catalytic machinery. The absence of mutations in (predicted) pockets and residues essential for heme binding indicates purifying selection acting to maintain the enzymatic core.

In addition to their structural similarities, CYP9Q3 and CYP336A1 exhibit complex epistatic architectures, in which adaptive effects arise from combinations of mutations rather than single substitutions (Ose et al., 2023) (Table S6). In CYP9Q3, the T62M variant displays both the highest mutational score (2.0) and the highest allele frequency (43.5%), yet occurs alone in only 21 individuals. This pattern implies that the structural or functional effects of T62M are context-dependent (a phenomenon known as epistasis), likely requiring compensatory mutations to maintain enzyme stability or modulate activity (Camps et al., 2007). Such interactions are common in P450 evolution, in which co-occurring mutations can mitigate deleterious impacts or synergistically enhance catalytic efficiency (Acevedo-Rocha et al., 2021; Ose et al., 2023).

Epistasis is even more pronounced in CYP336A1. A five-mutation haplotype (D202G, M207I, I222V, V226I, and Q238K) dominates population variation, with four of the five haplotypes never occurring in isolation (Table S6). These substitutions cluster around helix F′ and adjacent loop regions (Figure 9). This illustrates how detoxification enzymes evolve through coordinated networks of substitutions rather than isolated changes, as this enables rapid adaptation to chemical pressures while maintaining essential catalytic function (Camps et al., 2007; Ose et al., 2023; Pál et al., 2006). The presence of haplotypes with multiple SNPs in both genes underscores that adaptive potential resides not in individual variants but in their specific combinations and interactions.

### 4.5 Geographic and subspecies patterns in CYP diversity

Intracolonial genetic diversity is crucial for colony survival and fitness. It helps prevent infections, boosts growth, enhances productivity, and improves survival (Tarpy, 2003; Tarpy and Seeley, 2006Tarpy et al., 2013). CYP genes are highly polymorphic, a feature that helps populations adapt to dietary and environmental challenges. As a result, populations with greater CYP allelic diversity are expected to show increased resilience, as multiple enzyme variants can collectively detoxify a wider array of stressors (Mokhosoev et al., 2024).

Population differentiation was highly significant across all metrics: Hd, π, Tajima’s D, and F_ST_. This contrasts with housekeeping genes, which displayed no significant differentiation in any of these metrics. Thus, this confirms that the heterogeneous pattern observed in CYP reflects adaptive or demographic signals.

Our study provides the first population-level characterisation of CYP gene variation across *A. m.* subspecies and countries. Population differentiation is pronounced: Middle Eastern (Lebanon, Jordan, and UAE) populations consistently show high CYP diversity, whereas Iberian (Portugal and Spain) populations exhibit low diversity. This gradient was strongly driven by CYP3 (Hd = 50.67, p < 0.001), with Portugal differing significantly from Jordan, Malta, the UAE, and Slovenia in CYP-3-specific post-hoc tests.

Omani bees exhibited unique haplotypes, suggesting strong geographic isolation or intense local selection. The low diversity in Iberian populations contrasts sharply with that in North African populations, despite their geographic proximity.

Subspecies-level variation was equally striking: *A. m. siciliana* and *A. m. sahariensis* retain high CYP diversity across most clans, while subspecies like *A. m. mellifera* show severely reduced variation. The observed high diversity in African honey bees was expected, as genetic variability in these subspecies is generally high due to large effective population (propelled by the absecence of glaciation in Africa, as well as their migratory behaviour and swarm tendencies) and high mating frequency (Estoup et al., 1995; Franck et al., 2001).

However, it is important to note that all *A. m. mellifera* samples considered pure in our analysis originated from French conservation populations (mostly from Porquerolles). Wragg et al. (2022) highlighted the crucial role of such conservatories in safeguarding the genetic background of *A. m. mellifera*, which is threatened by C-lineage introgression due to modern beekeeping practices..

F_ST_ patterns offer strong support for population-specific adaptation. The repeated identification of CYP3 genes involved in xenobiotic detoxification, showing high differentiation across both countries and subspecies, suggets that these loci are key drivers of population structure, potentially under divergent selection pressures from regional flora, environmental conditions, or pesticide exposure histories. This is confirmed by the clan enrichment analysis of F_ST_ ≥ 0.9 variants: CYP3 genes were significantly enriched in both subspecies and country analyses, whereas CYP2 was significantly depleted. CYP3 accounted for 64.7% and 67.1% of all high F_ST_ variants in the subspecies and country analyses, respectively. This combined with the MK evidence of positive selection in CYP3 genes suggests that this clan is part of the primary adaptive interface between *A. mellifera* and its environment.

### 4.6 Implications for pesticide risk assessment and resistance management

Evidence suggests that, as in humans, a few CYPs bear the bulk of the detoxification load, while others play auxiliary or redundant roles (Haas et al., 2021; Manjon et al., 2018).

Current regulatory frameworks largely assume genetic homogeneity within *A. mellifera* and evaluate toxicity using limited laboratory subspecies (EFSA et al., 2023). Our comprehensive analysis demonstrates substantial and geographically structured CYP diversity across subspecies and geographic populations.

This genetic heterogeneity fundamentally challenges the current ’one-size-fits-all’ approach to risk assessment. Incorporating genetic variation into ecotoxicological and risk assessment models represents a critical and necessary advancement. By identifying populations with low detoxification gene diversity, regulators could implement region-specific restrictions or application guidelines, thereby establishing pharmacogenomic-informed beekeeping areas. Conversely, understanding which haplotypes confer metabolic resistance could guide breeding programs aimed at enhancing colony resilience without relying solely on reduced pesticide exposure, a challenging goal given agricultural intensification.

Just as monitoring insecticide resistance alleles in agricultural pests informs management strategies (Ffrench-Constant et al., 2004), tracking CYP haplotype frequencies for key genes across honey bee populations could help to detect emerging resistance or identify vulnerable populations. Rare haplotypes that increase rapidly in frequency may indicate ongoing selection from pesticide exposure, providing early warning signals for ecological monitoring programs. Haplotype analyses are especially informative for this highly polymorphic CYP genes.

Notably, we detected a previously reported CYP9Q3 haplotype (haplotype L – T62M, M288T, V380A) associated with reduced survival under clothianidin exposure (Tsvetkov et al., 2023) in 14 individuals across Cyprus (16%), Jordan (1.7%), and Turkey (2.2%). Its low frequency may reflect selective pressure from neonicotinoid exposure, illustrating that monitoring allele and haplotype frequencies can help identify populations at increased risk.

The geographic patterns observed could also enable the retrospective assessment of pesticide impacts. If historical pesticide use differs between regions showing high versus low CYP diversity, this natural variation could reveal long-term evolutionary consequences of chemical exposure.

### 4.7 Conservation implications and future directions

The pronounced diversity differences between subspecies have important conservation implications. *A. m. sahariensis* and *A. m. siciliana* show substantial CYP allelic diversity that may prove critical as climate change and agricultural intensification alter environmental pressures. The concentration of unique haplotypes in North African and Middle Eastern subspecies suggests that these populations contain adaptive potential crucial for long-term species resilience.

The missense haplotypes in high-diversity genes, particularly CYP9Q3 and CYP336A1, provide targets for functional validation studies. Characterising the substrate specificities and catalytic efficiencies of major haplotypes could reveal how genetic variation translates into phenotypic differences in pesticide tolerance. Heterologous expression systems combined with molecular docking and molecular dynamics simulations can predict functional consequences before costly *in vivo* validation.

The catalogue we provide sets the stage for phenotype–genotype association studies. Controlled exposure experiments comparing pesticide metabolism rates or colony survival across known haplotypes could definitively link genetic variation to functional outcomes. Such studies would validate *in silico* predictions, enabling predictive models for population-level responses. Integration with transcriptomic data would further enhance utility: expression differences between haplotypes or differential inducibility following xenobiotic exposure could explain why some variants confer protection despite modest amino acid changes. CYP genes showing both high genetic diversity and expression plasticity may be particularly important for adaptive responses.

### 4.8 Study limitations and broader perspectives

Although this study provides the first broad survey of CYP variation across *A. mellifera* populations, several considerations merit discussion. First, the reference genome bias, evident in genes such as CYP6AS18, where reference alleles are completely absent, suggests caution is needed when interpreting "wild-type" function based on genomic references derived from limited samples.

Second, by focusing exclusively on SNPs, we may have missed other forms of genetic diversity that influence CYP function. Structural variants (copy number variations, insertions/deletions), regulatory variants affecting expression levels, and epigenetic modifications are known to influence CYP function in other organisms (Ingelman-Sundberg et al., 2007) but remain largely unexplored in honey bees. Future studies incorporating whole-genome sequencing and RNA-seq data would provide a more complete picture of detoxification gene diversity.

Expanding this framework to other detoxification gene families (carboxyl/cholinesterases (CCEs), glutathione S-transferases (GSTs), ATP-binding cassette transporters (ABCs), and uridine 5′-diphospho-glucuronosyltransferases (UGTs)) would create a comprehensive pharmacogenomic resource for honey bees. Cross-family analyses might reveal co-adapted gene complexes or identify populations with globally enhanced or diminished detoxification capacity. The interplay between different detoxification pathways (phase I metabolism by CYPs, phase II conjugation by GSTs and UGTs, and phase III transport by ABCs) determines overall xenobiotic tolerance, and understanding genetic variation across this entire network would provide the most complete picture of adaptive potential.

As honey bee colony losses continue globally (Potts et al., 2010), resources such as this catalogue are increasingly urgent for evidence-based conservation and sustainable apiculture. By bridging genomics, evolutionary biology, and toxicology, we hope that this work will catalyse further research into the genetic basis of pollinator resilience and inform management strategies that preserve both biodiversity and ecosystem services.

---

**Figure S1.**
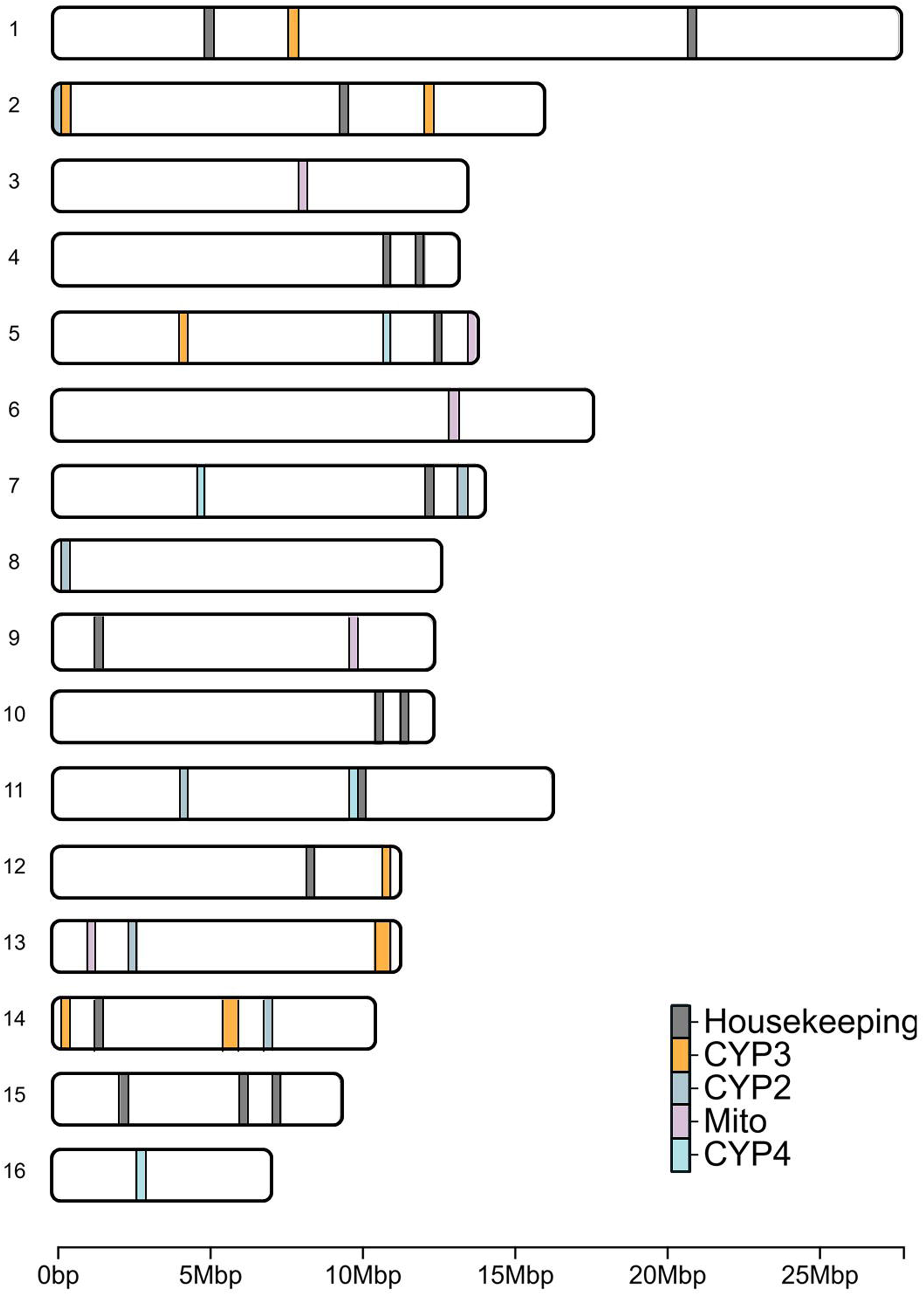
Distribution of CYP (grouped by clan) and housekeeping genes across the 16 chromosomes of the honey bee (Apis mellifera). Mito – Mitochondrial CYP clan

**Figure S2.**
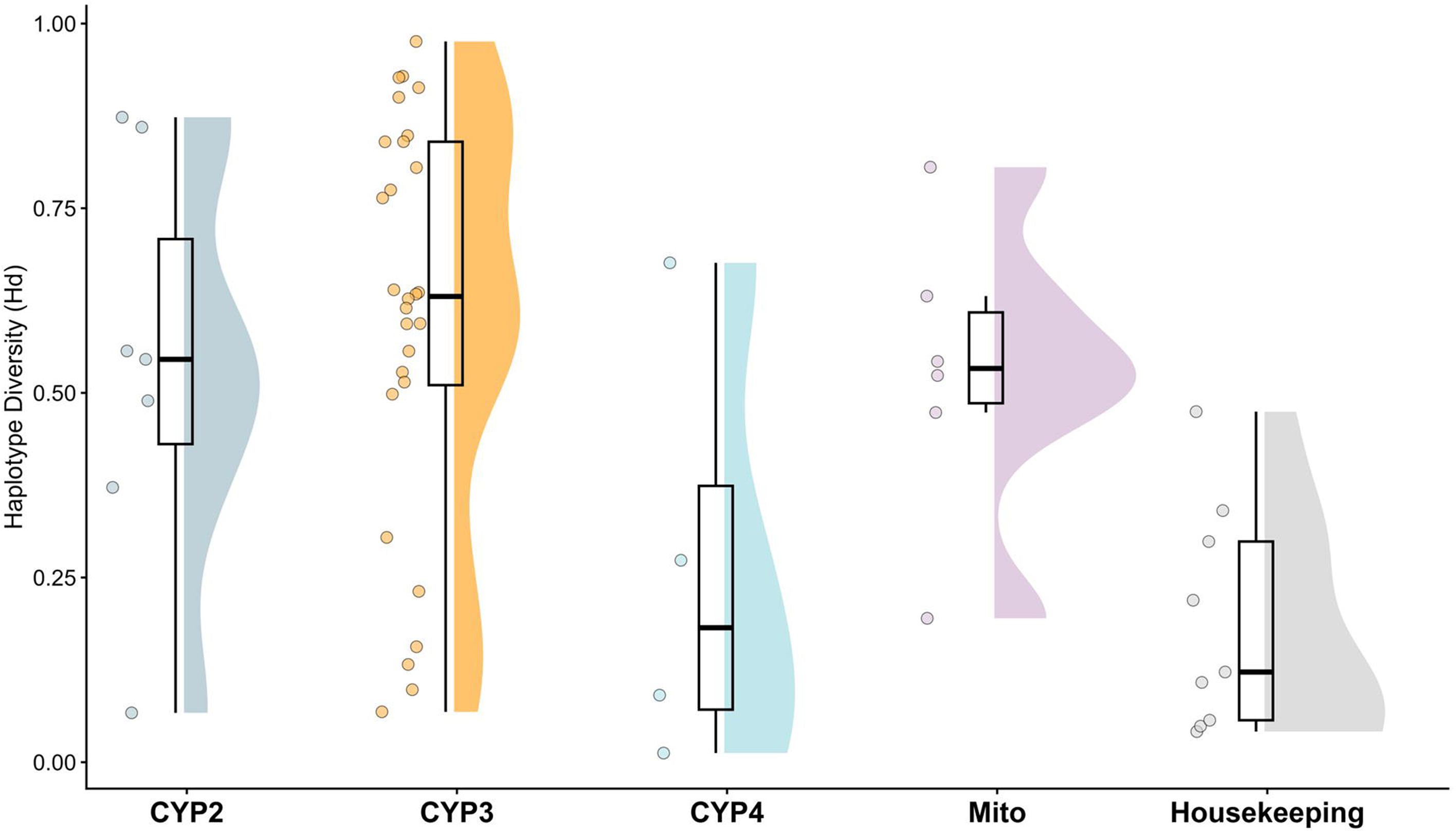
Haplotype diversity across CYP and housekeeping genes. The CYP3 clan showed extreme diversity variation, containing both the most diverse gene (CYP9S1; Hd = 0.976) and one of the least diverse (CYP6AS8; Hd = 0.068).

**Figure S3.**
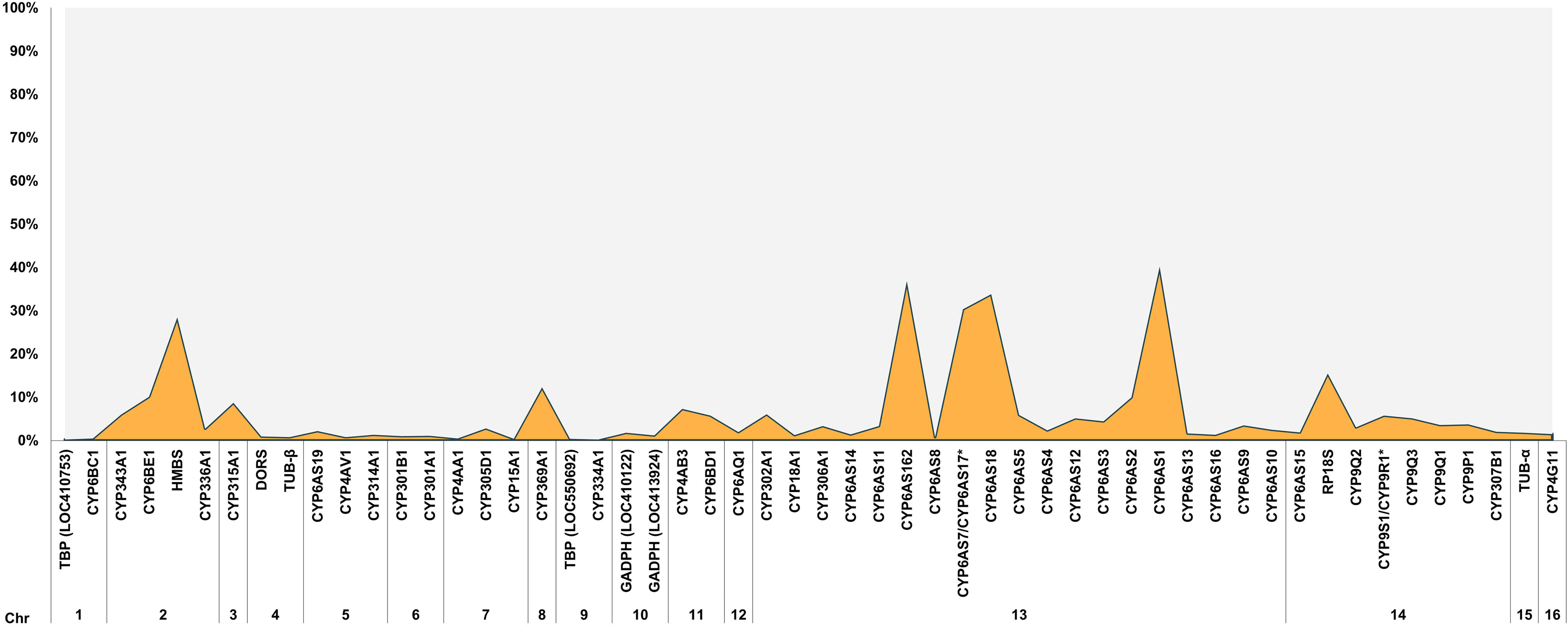
Relative distribution of missing data per gene.

**Figure S4.**
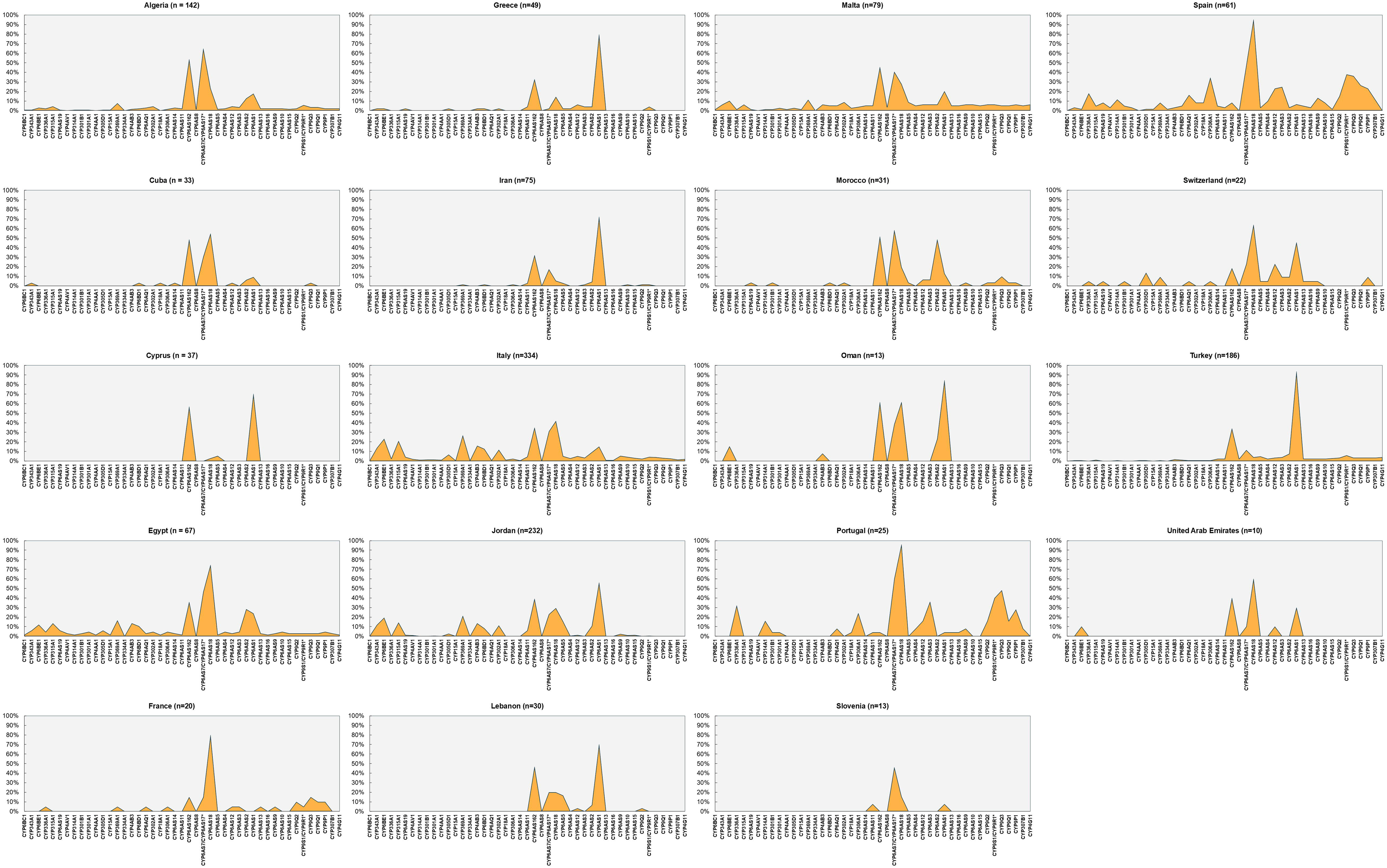
Relative distribution of missing data across countries.

**Figure S5.**
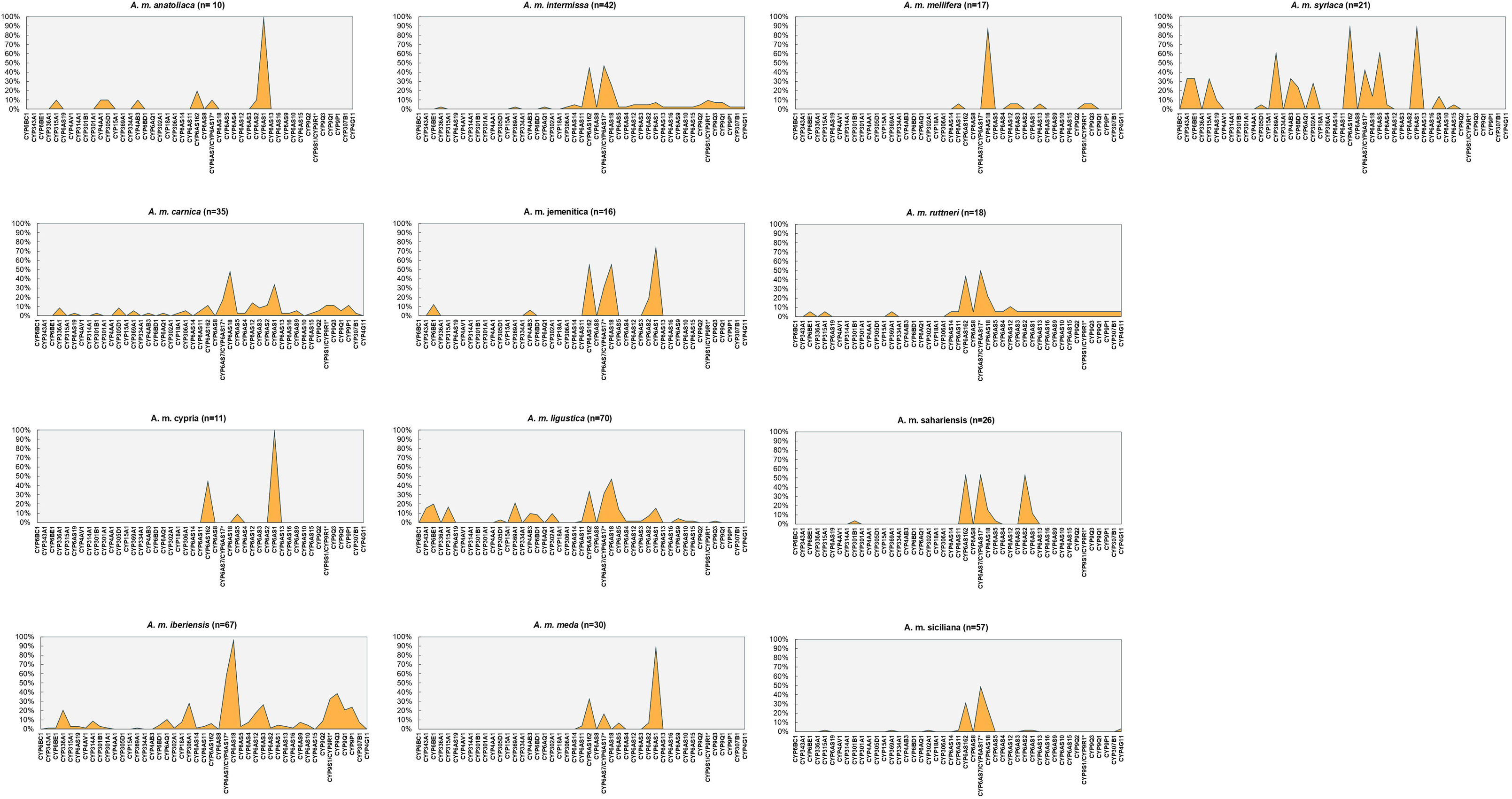
Relative distribution of missing data per CYP genes across honey bee subspecies.

**Figure S6.**
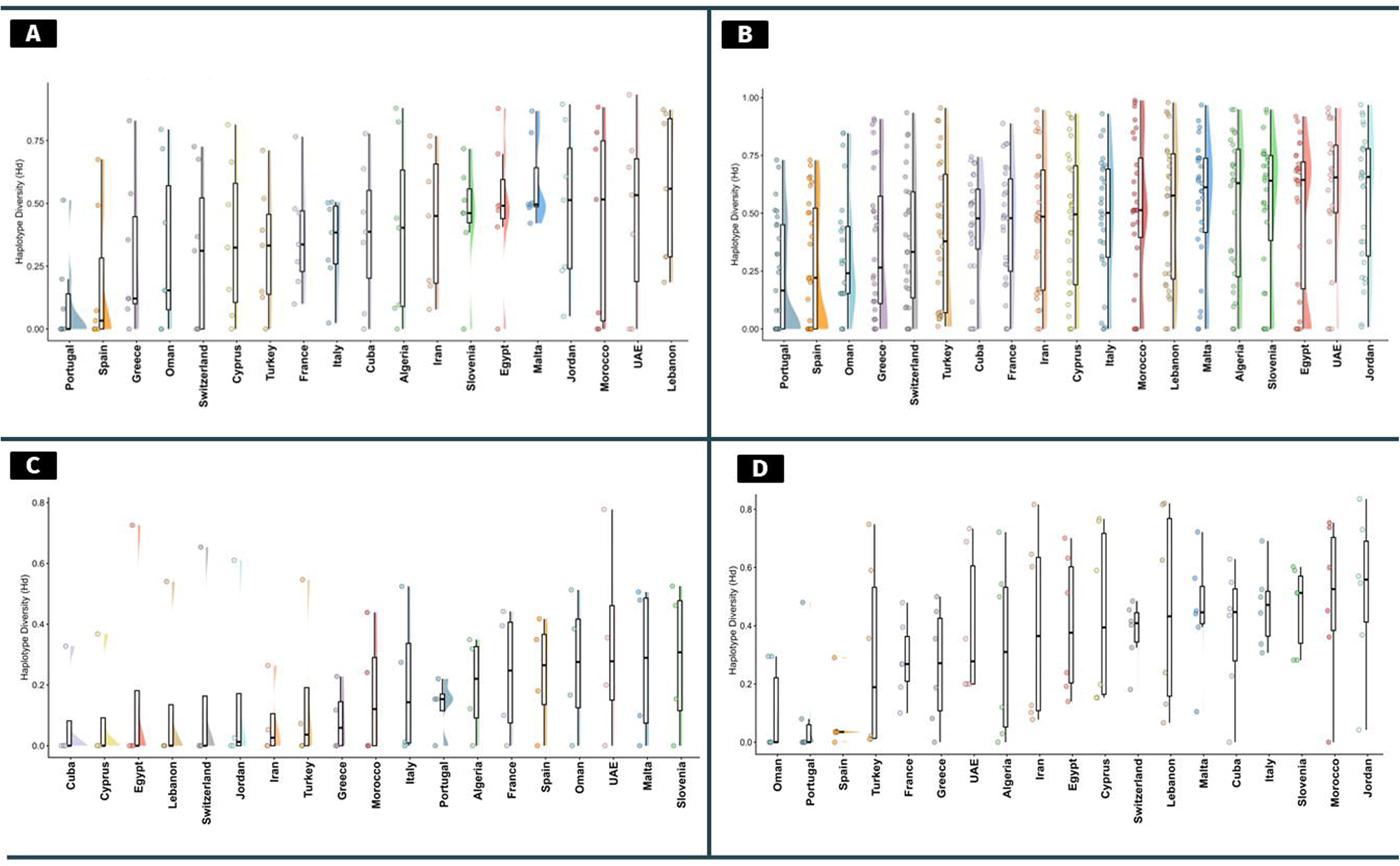
Haplotype Diversity (Hd) of Honey Bees Across Countries. Haplotype diversity values are categorized by clan: A) CYP2 clan, B) CYP3 clan, C) CYP4 clan, and D) Mitochondrial clan

**Figure S7.**
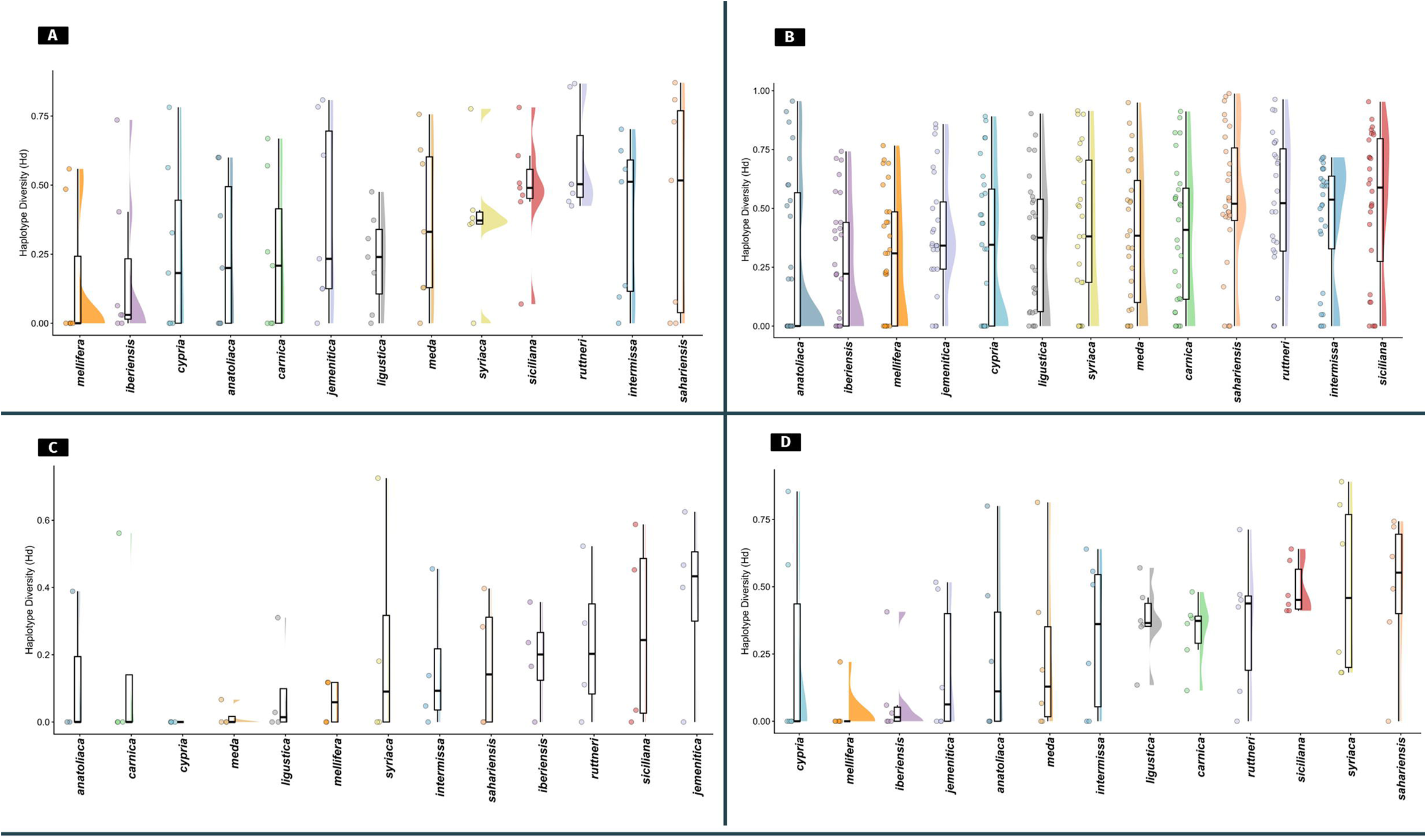
Haplotype diversity (Hd) across honey bee subspecies. Haplotype diversity values are categorized by clan: A – CYP2 clan, B – CYP3 clan, C – CYP4 clan, and D – Mitochondrial clan.

## References

Acevedo-Rocha, C.G., Li, A., D’Amore, L., Hoebenreich, S., Sanchis, J., Lubrano, P., Ferla, M.P., Garcia-Borràs, M., Osuna, S., Reetz, M.T., 2021. Pervasive cooperative mutational effects on multiple catalytic enzyme traits emerge via long-range conformational dynamics. Nat. Commun. 12, 1621. 10.1038/s41467-021-21833-w

Arena, M., Sgolastra, F., 2014. A meta-analysis comparing the sensitivity of bees to pesticides. Ecotoxicology 23, 324–334. 10.1007/s10646-014-1190-1

Aupinel, P., Fortini, D., Michaud, B., Medrzycki, P., Padovani, E., Przygoda, D., Maus, C., Charriere, J.-D., Kilchenmann, V., Riessberger-Galle, U., 2010. Honey bee brood ring-test: method for testing pesticide toxicity on honeybee brood in laboratory conditions. Julius-Kühn-Archiv 96.

Bass, C., Hayward, A., Troczka, B.J., Haas, J., Nauen, R., 2024. The molecular determinants of pesticide sensitivity in bee pollinators. Science of The Total Environment 915, 170174. 10.1016/J.SCITOTENV.2024.170174

Bateman, A., Martin, M.J., Orchard, S., Magrane, M., Adesina, A., Ahmad, S., Bowler-Barnett, E.H., Bye-A-Jee, H., Carpentier, D., Denny, P., Fan, J., Garmiri, P., da Costa Gonzales, L.J., Hussein, A., Ignatchenko, A., Insana, G., Ishtiaq, R., Joshi, V., Jyothi, D., Kandasaamy, S., Lock, A., Luciani, A., Luo, J., Lussi, Y., Marin, J.S.M., Raposo, P., Rice, D.L., Santos, R., Speretta, E., Stephenson, J., Totoo, P., Tyagi, N., Urakova, N., Vasudev, P., Warner, K., Wijerathne, S., Yu, C.W.H., Zaru, R., Bridge, A.J., Aimo, L., Argoud-Puy, G., Auchincloss, A.H., Axelsen, K.B., Bansal, P., Baratin, D., Batista Neto, T.M., Blatter, M.C., Bolleman, J.T., Boutet, E., Breuza, L., Gil, B.C., Casals-Casas, C., Echioukh, K.C., Coudert, E., Cuche, B., de Castro, E., Estreicher, A., Famiglietti, M.L., Feuermann, M., Gasteiger, E., Gaudet, P., Gehant, S., Gerritsen, V., Gos, A., Gruaz, N., Hulo, C., Hyka-Nouspikel, N., Jungo, F., Kerhornou, A., Le Mercier, P., Lieberherr, D., Masson, P., Morgat, A., Paesano, S., Pedruzzi, I., Pilbout, S., Pourcel, L., Poux, S., Pozzato, M., Pruess, M., Redaschi, N., Rivoire, C., Sigrist, C.J.A., Sonesson, K., Sundaram, S., Sveshnikova, A., Wu, C.H., Arighi, C.N., Chen, C., Chen, Y., Huang, H., Laiho, K., Lehvaslaiho, M., McGarvey, P., Natale, D.A., Ross, K., Vinayaka, C.R., Wang, Y., Zhang, J., 2025. UniProt: the Universal Protein Knowledgebase in 2025. Nucleic Acids Res. 53, D609–D617. 10.1093/NAR/GKAE1010

Berenbaum, M.R., 2002. Postgenomic chemical ecology: from genetic code to ecological interactions. J. Chem. Ecol. 28, 873–896. 10.1023/A:1015260931034

Blacquière, T., Smagghe, G., van Gestel, C.A.M., Mommaerts, V., 2012. Neonicotinoids in bees: a review on concentrations, side-effects and risk assessment. Ecotoxicology 21, 973–992. 10.1007/s10646-012-0863-x

Bogwitz, M.R., Chung, H., Magoc, L., Rigby, S., Wong, W., O’Keefe, M., McKenzie, J.A., Batterham, P., Daborn, P.J., 2005. Cyp12a4 confers lufenuron resistance in a natural population of Drosophila melanogaster. Proc. Natl. Acad. Sci. U. S. A. 102, 12807–12812. 10.1073/pnas.0503709102

Boncristiani, H., Underwood, R., Schwarz, R., Evans, J.D., Pettis, J., Vanengelsdorp, D., 2012. Direct effect of acaricides on pathogen loads and gene expression levels in honey bees Apis mellifera. J. Insect Physiol. 58, 613–620. 10.1016/J.JINSPHYS.2011.12.011

Campos, M., Markham, K.R., Mitchell, K.A., da Cunha, A.P., 1997. An approach to the characterization of bee pollens via their flavonoid/phenolic profiles. Phytochemical Analysis 8, 181–185. 10.1002/(SICI)1099-1565(199707)8:4<181::AID-PCA359>3.0.CO;2-A

Camps, M., Herman, A., Loh, E., Loeb, L.A., 2007. Genetic Constraints on Protein Evolution. Crit. Rev. Biochem. Mol. Biol. 42, 313–326. 10.1080/10409230701597642

Capriotti, E., Fariselli, P., Casadio, R., 2005. I-Mutant2.0: Predicting stability changes upon mutation from the protein sequence or structure. Nucleic Acids Res. 33. 10.1093/nar/gki375

Chaimanee, V., Evans, J.D., Chen, Y., Jackson, C., Pettis, J.S., 2016. Sperm viability and gene expression in honey bee queens (Apis mellifera) following exposure to the neonicotinoid insecticide imidacloprid and the organophosphate acaricide coumaphos. J. Insect Physiol. 89, 1–8. 10.1016/J.JINSPHYS.2016.03.004

Claudianos, C., Ranson, H., Johnson, R.M., Biswas, S., Schuler, M.A., Berenbaum, M.R., Feyereisen, R., Oakeshott, J.G., 2006. A deficit of detoxification enzymes: pesticide sensitivity and environmental response in the honeybee. Insect Mol. Biol. 15, 615. 10.1111/J.1365-2583.2006.00672.X

Daborn, P.J., Lumb, C., Boey, A., Wong, W., ffrench-Constant, R.H., Batterham, P., 2007. Evaluating the insecticide resistance potential of eight Drosophila melanogaster cytochrome P450 genes by transgenic over-expression. Insect Biochem. Mol. Biol. 37, 512–519. 10.1016/j.ibmb.2007.02.008

Danka, R.G., Rinderer, T.E., Hellmich, R.L., Collins, A.M., 1986. Comparative Toxicities of Four Topically Applied Insecticides to Africanized and European Honey Bees (Hymenoptera: Apidae). J. Econ. Entomol. 79, 18–21. 10.1093/jee/79.1.18

Derecka, K., Blythe, M.J., Malla, S., Genereux, D.P., Guffanti, A., Pavan, P., Moles, A., Snart, C., Ryder, T., Ortori, C.A., Barrett, D.A., Schuster, E., Stöger, R., 2013. Transient Exposure to Low Levels of Insecticide Affects Metabolic Networks of Honeybee Larvae. PLoS One 8, e68191. 10.1371/JOURNAL.PONE.0068191

Dermauw, W., Van Leeuwen, T., Feyereisen, R., 2020. Diversity and evolution of the P450 family in arthropods. Insect Biochem. Mol. Biol. 127, 103490. 10.1016/J.IBMB.2020.103490

Dyer, S.C., Austine-Orimoloye, O., Azov, A.G., Barba, M., Barnes, I., Barrera-Enriquez, V.P., Becker, A., Bennett, R., Beracochea, M., Berry, A., Bhai, J., Bhurji, S.K., Boddu, S., Branco Lins, P.R., Brooks, L., Ramaraju, S.B., Campbell, L.I., Martinez, M.C., Charkhchi, M., Cortes, L.A., Davidson, C., Denni, S., Dodiya, K., Donaldson, S., Houdaigui, B. El, Naboulsi, T. El, Falola, O., Fatima, R., Genez, T., Martinez, J.G., Gurbich, T., Hardy, M., Hollis, Z., Hunt, T., Kay, M., Kaykala, V., Lemos, D., Lodha, D., Mathlouthi, N., Merino, G.A., Merritt, R., Mirabueno, L.P., Mushtaq, A., Hossain, S.N., Pérez-Silva, J.G., Perry, M., Piližota, I., Poppleton, D., Prosovetskaia, I., Raj, S., Salam, A.I.A., Saraf, S., Saraiva-Agostinho, N., Sinha, S., Sipos, B., Sitnik, V., Steed, E., Suner, M.M., Surapaneni, L., Sutinen, K., Tricomi, F.F., Tsang, I., Urbina-Gómez, D., Veidenberg, A., Walsh, T.A., Willhoft, N.L., Allen, J., Alvarez-Jarreta, J., Chakiachvili, M., Cheema, J., da Rocha, J.B., De Silva, N.H., Giorgetti, S., Haggerty, L., Ilsley, G.R., Keatley, J., Loveland, J.E., Moore, B., Mudge, J.M., Naamati, G., Tate, J., Trevanion, S.J., Winterbottom, A., Flint, B., Frankish, A., Hunt, S.E., Finn, R.D., Freeberg, M.A., Harrison, P.W., Martin, F.J., Yates, A.D., 2025. Ensembl 2025. Nucleic Acids Res. 53, D948–D957. 10.1093/NAR/GKAE1071

Eddy, S.R., 2004. Where did the BLOSUM62 alignment score matrix come from? Nat. Biotechnol. 22, 1035–1036. 10.1038/NBT0804-1035;

EFSA, E.F.S.A., Adriaanse, P., Arce, A., Focks, A., Ingels, B., Jölli, D., Lambin, S., Rundlöf, M., Süßenbach, D., Del Aguila, M., Ercolano, V., Ferilli, F., Ippolito, A., Szentes, C., Neri, F.M., Padovani, L., Rortais, A., Wassenberg, J., Auteri, D., 2023. Revised guidance on the risk assessment of plant protection products on bees (Apis mellifera, Bombus spp. and solitary bees). EFSA Journal 21, e07989. 10.2903/J.EFSA.2023.7989;

Ehrt, C., Schulze, T., Graef, J., Diedrich, K., Pletzer-Zelgert, J., Rarey, M., 2025. ProteinsPlus: a publicly available resource for protein structure mining. Nucleic Acids Res. 53, W478–W484. 10.1093/NAR/GKAF377

Eilbeck, K., Lewis, S.E., Mungall, C.J., Yandell, M., Stein, L., Durbin, R., Ashburner, M., 2005. The Sequence Ontology: a tool for the unification of genome annotations. Genome Biol. 6, R44. 10.1186/GB-2005-6-5-R44

El Agrebi, N., De Smet, L., Douny, C., Scippo, M.L., Svečnjak, L., de Graaf, D.C., Saegerman, C., 2024. A field realistic model to assess the effects of pesticides residues and adulterants on honey bee gene expression. PLoS One 19, e0302183. 10.1371/JOURNAL.PONE.0302183

Elzen, P.J., Elzen, G.W., Rubink, W., 2003. Comparative susceptibility of European and Africanized honey bee (Hymenoptera: Apidae) ecotypes to several insecticide classes. Southwest Entomol 28, 255–260.

Emms, D.M., Kelly, S., 2019. OrthoFinder: phylogenetic orthology inference for comparative genomics. Genome Biol. 20, 238. 10.1186/s13059-019-1832-y

Estoup, A., Garnery, L., Solignac, M., Cornuet, J.M., 1995. Microsatellite variation in honey bee (Apis mellifera L.) populations: hierarchical genetic structure and test of the infinite allele and stepwise mutation models. Genetics 140, 679–695. 10.1093/genetics/140.2.679

Feyereisen, R., 2012. Insect CYP Genes and P450 Enzymes, in: Gilbert, L.I. (Ed.), Insect Molecular Biology and Biochemistry. Academic Press, pp. 236–316. 10.1016/B978-0-12-384747-8.10008-X

Feyereisen, R., 2006. Evolution of insect P450. Biochem. Soc. Trans. 34, 1252–1255. 10.1042/BST0341252

Ffrench-Constant, R.H., Daborn, P.J., Le Goff, G., 2004. The genetics and genomics of insecticide resistance. Trends in Genetics 20, 163–170. 10.1016/J.TIG.2004.01.003

Franck, P., Garnery, L., Loiseau, A., Oldroyd, B.P., Hepburn, H.R., Solignac, M., Cornuet, J.M., 2001. Genetic diversity of the honeybee in Africa: microsatellite and mitochondrial data. Heredity 2001 86:4 86, 420–430. 10.1046/j.1365-2540.2001.00842.x

Frenz, B., Lewis, S.M., King, I., DiMaio, F., Park, H., Song, Y., 2020. Prediction of Protein Mutational Free Energy: Benchmark and Sampling Improvements Increase Classification Accuracy. Front. Bioeng. Biotechnol. 8, 558247. 10.3389/FBIOE.2020.558247/FULL

Fujita, T., Kozuka-Hata, H., Hori, Y., Takeuchi, J., Kubo, T., Oyama, M., 2018. Shotgun proteomics deciphered age/division of labor-related functional specification of three honeybee (Apis mellifera L.) exocrine glands. PLoS One 13, e0191344. 10.1371/JOURNAL.PONE.0191344

Good, R.T., Gramzow, L., Battlay, P., Sztal, T., Batterham, P., Robin, C., 2014. The Molecular Evolution of Cytochrome P450 Genes within and between Drosophila Species. Genome Biol. Evol. 6, 1118–1134. 10.1093/gbe/evu083

Goulson, D., Nicholls, E., Botías, C., Rotheray, E.L., 2015. Bee declines driven by combined Stress from parasites, pesticides, and lack of flowers. Science (1979). 347. 10.1126/science.1255957

Gregorc, A., Silva-Zacarin, E.C.M., Carvalho, S.M., Kramberger, D., Teixeira, E.W., Malaspina, O., 2016. Effects of Nosema ceranae and thiametoxam in Apis mellifera: A comparative study in Africanized and Carniolan honey bees. Chemosphere 147, 328–336. 10.1016/j.chemosphere.2015.12.030

Guzov, V.M., Unnithan, G.C., Chernogolov, A.A., Feyereisen, R., 1998. CYP12A1, a mitochondrial cytochrome P450 from the house fly. Arch. Biochem. Biophys. 359, 231–240. 10.1006/abbi.1998.0901

Gyulkhandanyan, A., Rezaie, A.R., Roumenina, L., Lagarde, N., Fremeaux-Bacchi, V., Miteva, M.A., Villoutreix, B.O., 2020. Analysis of protein missense alterations by combining sequence- and structure-based methods. Mol. Genet. Genomic Med. 8, e1166. 10.1002/MGG3.1166

Haas, J., Beck, E., Troczka, B.J., Hayward, A., Hertlein, G., Zaworra, M., Lueke, B., Buer, B., Maiwald, F., Beck, M.E., Nebelsiek, B., Glaubitz, J., Bass, C., Nauen, R., 2023. A conserved hymenopteran-specific family of cytochrome P450s protects bee pollinators from toxic nectar alkaloids. Sci. Adv. 9. 10.1126/sciadv.adg0885

Haas, J., Glaubitz, J., Koenig, U., Nauen, R., 2022a. A mechanism-based approach unveils metabolic routes potentially mediating chlorantraniliprole synergism in honey bees, Apis mellifera L., by azole fungicides. Pest Manag. Sci. 78, 965–973. 10.1002/PS.6706

Haas, J., Hayward, A., Buer, B., Maiwald, F., Nebelsiek, B., Glaubitz, J., Bass, C., Nauen, R., 2022b. Phylogenomic and functional characterization of an evolutionary conserved cytochrome P450-based insecticide detoxification mechanism in bees. Proceedings of the National Academy of Sciences 119. 10.1073/pnas.2205850119

Haas, J., Zaworra, M., Glaubitz, J., Hertlein, G., Kohler, M., Lagojda, A., Lueke, B., Maus, C., Almanza, M.T., Davies, T.G.E., Bass, C., Nauen, R., 2021. A toxicogenomics approach reveals characteristics supporting the honey bee (Apis mellifera L.) safety profile of the butenolide insecticide flupyradifurone. Ecotoxicol. Environ. Saf. 217, 112247. 10.1016/J.ECOENV.2021.112247

Haberle, V., Lenhard, B., 2016. Promoter architectures and developmental gene regulation. Semin. Cell Dev. Biol. 57, 11–23. 10.1016/J.SEMCDB.2016.01.014

Han, H., Yang, Y., Hu, J., Wang, Y., Zhao, Z., Ma, R., Gao, L., Guo, Y., 2022. Identification and Characterization of CYP6 Family Genes from the Oriental Fruit Moth (Grapholita molesta) and Their Responses to Insecticides. Insects 13, 300. 10.3390/INSECTS13030300/S1

Harris, A.M., DeGiorgio, M., 2017. An unbiased estimator of gene diversity with improved variance for samples containing related and inbred individuals of any ploidy. G3: Genes, Genomes, Genetics 7, 671–691. 10.1534/G3.116.037168/-/DC1

Hayward, A., Beadle, K., Singh, K.S., Exeler, N., Zaworra, M., Almanza, M.-T., Nikolakis, A., Garside, C., Glaubitz, J., Bass, C., Nauen, R., 2019. The leafcutter bee, Megachile rotundata, is more sensitive to N-cyanoamidine neonicotinoid and butenolide insecticides than other managed bees. Nat. Ecol. Evol. 3, 1521–1524. 10.1038/s41559-019-1011-2

Hayward, A., Hunt, B.J., Haas, J., Bushnell-Crowther, E., Troczka, B.J., Pym, A., Beadle, K., Field, J., Nelson, D.R., Nauen, R., Bass, C., 2023. A cytochrome P450 insecticide detoxification mechanism is not conserved across the Megachilidae family of bees. Evol. Appl. 17, e13625. 10.1111/EVA.13625

Hayward, A.J., O’Reilly, A.O., Nauen, R., Bass, C., Troczka, B.J., 2026. An in silico pipeline for enzyme-substrate modelling using arthropod P450s. Pestic. Biochem. Physiol. 216, 106816. 10.1016/J.PESTBP.2025.106816

Healy, J., McInnes, L., 2024. Uniform manifold approximation and projection. Nature Reviews Methods Primers 4, 1–15. 10.1038/S43586-024-00363-X

Henikoff, S., Henikoff, J.G., 1992. Amino acid substitution matrices from protein blocks. Proc. Natl. Acad. Sci. U. S. A. 89, 10915. 10.1073/PNAS.89.22.10915

Henriques, D., Wallberg, A., Chávez-Galarza, J., Johnston, J.S., Webster, M.T., Pinto, M.A., 2018. Whole genome SNP-associated signatures of local adaptation in honeybees of the Iberian Peninsula. Sci. Rep. 8, 11145. 10.1038/S41598-018-29469-5

Ingelman-Sundberg, M., Sim, S.C., Gomez, A., Rodriguez-Antona, C., 2007. Influence of cytochrome P450 polymorphisms on drug therapies: Pharmacogenetic, pharmacoepigenetic and clinical aspects. Pharmacol. Ther. 116, 496–526. 10.1016/j.pharmthera.2007.09.004

Ittisoponpisan, S., Islam, S.A., Khanna, T., Alhuzimi, E., David, A., Sternberg, M.J.E., 2019. Can Predicted Protein 3D Structures Provide Reliable Insights into whether Missense Variants Are Disease Associated? J. Mol. Biol. 431, 2197–2212. 10.1016/J.JMB.2019.04.009

Iwasa, T., Motoyama, N., Ambrose, J.T., Roe, R.M., 2004. Mechanism for the differential toxicity of neonicotinoid insecticides in the honey bee, Apis mellifera. Crop Protection 23, 371–378. 10.1016/j.cropro.2003.08.018

Klein, A.M., Vaissière, B.E., Cane, J.H., Steffan-Dewenter, I., Cunningham, S.A., Kremen, C., Tscharntke, T., 2007. Importance of pollinators in changing landscapes for world crops. Proceedings of the Royal Society B: Biological Sciences 274, 303–313. 10.1098/rspb.2006.3721

Kopelman, N.M., Mayzel, J., Jakobsson, M., Rosenberg, N.A., Mayrose, I., 2015. Clumpak: a program for identifying clustering modes and packaging population structure inferences across K. Mol. Ecol. Resour. 15, 1179–1191. 10.1111/1755-0998.12387

Laurino, D., Manino, A., Patetta, A., Porporato, M., 2013. Toxicity of neonicotinoid insecticides on different honey bee genotypes. Bull. Insectology 66, 119–126.

Li, H., Liu, S., Chen, L., Luo, J., Zeng, D., Li, X., 2021. Juvenile hormone and transcriptional changes in honey bee worker larvae when exposed to sublethal concentrations of thiamethoxam. Ecotoxicol. Environ. Saf. 225, 112744. 10.1016/J.ECOENV.2021.112744

Li, X., Schuler, M.A., Berenbaum, M.R., 2007a. Molecular Mechanisms of Metabolic Resistance to Synthetic and Natural Xenobiotics. Annu. Rev. Entomol. 52, 231–253. 10.1146/annurev.ento.51.110104.151104

Li, X., Schuler, M.A., Berenbaum, M.R., 2007b. Molecular mechanisms of metabolic resistance to synthetic and natural xenobiotics. Annu. Rev. Entomol. 52, 231–253. 10.1146/ANNUREV.ENTO.51.110104.151104/CITE/REFWORKS

Lü, J., Yang, C., Zhang, Y., Pan, H., 2018. Selection of reference genes for the normalization of RT-qPCR data in gene expression studies in insects: A systematic review. Front. Physiol. 9, 418953. 10.3389/FPHYS.2018.01560

Manjon, C., Troczka, B.J., Zaworra, M., Beadle, K., Randall, E., Hertlein, G., Singh, K.S., Zimmer, C.T., Homem, R.A., Lueke, B., Reid, R., Kor, L., Kohler, M., Benting, J., Williamson, M.S., Davies, T.G.E., Field, L.M., Bass, C., Nauen, R., 2018. Unravelling the Molecular Determinants of Bee Sensitivity to Neonicotinoid Insecticides. Current Biology 28, 1137–1143.e5. 10.1016/j.cub.2018.02.045

Mao, W., Rupasinghe, S.G., Johnson, R.M., Zangerl, A.R., Schuler, M.A., Berenbaum, M.R., 2009. Quercetin-metabolizing CYP6AS enzymes of the pollinator Apis mellifera (Hymenoptera: Apidae). Comparative Biochemistry and Physiology - B Biochemistry and Molecular Biology 154, 427–434. 10.1016/j.cbpb.2009.08.008

Mao, W., Schuler, M.A., Berenbaum, M.R., 2011. CYP9Q-mediated detoxification of acaricides in the honey bee (Apis mellifera). Proc. Natl. Acad. Sci. U. S. A. 108, 12657–12662. 10.1073/PNAS.1109535108

McInnes, L., Healy, J., Melville, J., 2018. UMAP: Uniform Manifold Approximation and Projection for Dimension Reduction. ArXiv e-prints.

McLaren, W., Gil, L., Hunt, S.E., Riat, H.S., Ritchie, G.R.S., Thormann, A., Flicek, P., Cunningham, F., 2016. The Ensembl Variant Effect Predictor. Genome Biol. 17, 1–14. 10.1186/S13059-016-0974-4

Milone, J.P., Rinkevich, F.D., McAfee, A., Foster, L.J., Tarpy, D.R., 2020. Differences in larval pesticide tolerance and esterase activity across honey bee (Apis mellifera) stocks. Ecotoxicol. Environ. Saf. 206, 111213. 10.1016/j.ecoenv.2020.111213

Mokhosoev, I.M., Astakhov, D. V., Terentiev, A.A., Moldogazieva, N.T., 2024. Cytochrome P450 monooxygenase systems: Diversity and plasticity for adaptive stress response. Prog. Biophys. Mol. Biol. 193, 19–34. 10.1016/J.PBIOMOLBIO.2024.09.003

Müller, P., Donnelly, M.J., Ranson, H., 2007. Transcription profiling of a recently colonised pyrethroid resistant Anopheles gambiae strain from Ghana. BMC Genomics 2007 8:1 8, 36-. 10.1186/1471-2164-8-36

Murga-Moreno, J., Coronado-Zamora, M., Casillas, S., Barbadilla, A., 2022. impMKT: the imputed McDonald and Kreitman test, a straightforward correction that significantly increases the evidence of positive selection of the McDonald and Kreitman test at the gene level. G3: Genes|Genomes|Genetics 12, jkac206. 10.1093/g3journal/jkac206

Nauen, R., Bass, C., Feyereisen, R., Vontas, J., 2022. The Role of Cytochrome P450s in Insect Toxicology and Resistance. Annu. Rev. Entomol. 67, 105–124. 10.1146/annurev-ento-070621-061328

Nei, M., Li, W.H., 1979. Mathematical model for studying genetic variation in terms of restriction endonucleases. Proceedings of the National Academy of Sciences 76, 5269–5273. 10.1073/PNAS.76.10.5269

Nei, M., Roychoudhury, A.K., 1974. SAMPLING VARIANCES OF HETEROZYGOSITY AND GENETIC DISTANCE. Genetics 76, 379–390. 10.1093/GENETICS/76.2.379

Nei, M., Tajima, F., 1981. DNA POLYMORPHISM DETECTABLE BY RESTRICTION ENDONUCLEASES. Genetics 97, 145–163. 10.1093/GENETICS/97.1.145

OECD, 2017. Guidance on Exposure and Effects Testing for Assessing Risks to Bees.

Ose, N.J., Campitelli, P., Patel, R., Kumar, S., Ozkan, S.B., 2023. Protein dynamics provide mechanistic insights about epistasis among common missense polymorphisms. Biophys. J. 122, 2938–2947. 10.1016/j.bpj.2023.01.037

Pál, C., Papp, B., Lercher, M.J., 2006. An integrated view of protein evolution. Nat. Rev. Genet. 7, 337–348. 10.1038/nrg1838

Pisa, L., Goulson, D., Yang, E.C., Gibbons, D., Sánchez-Bayo, F., Mitchell, E., Aebi, A., van der Sluijs, J., MacQuarrie, C.J.K., Giorio, C., Long, E.Y., McField, M., Bijleveld van Lexmond, M., Bonmatin, J.M., 2017. An update of the Worldwide Integrated Assessment (WIA) on systemic insecticides. Part 2: impacts on organisms and ecosystems. Environ. Sci. Pollut. Res. Int. 28, 11749. 10.1007/s11356-017-0341-3

Potts, S.G., Biesmeijer, J.C., Kremen, C., Neumann, P., Schweiger, O., Kunin, W.E., 2010. Global pollinator declines: trends, impacts and drivers. Trends Ecol. Evol. 25, 345–353. 10.1016/J.TREE.2010.01.007

Purcell, S., Neale, B., Todd-Brown, K., Thomas, L., Ferreira, M.A.R., Bender, D., Maller, J., Sklar, P., De Bakker, P.I.W., Daly, M.J., Sham, P.C., 2007. PLINK: A tool set for whole-genome association and population-based linkage analyses. Am. J. Hum. Genet. 81, 559–575. 10.1086/519795

Qiu, Y., Tittiger, C., Wicker-Thomas, C., Le Goff, G., Young, S., Wajnberg, E., Fricaux, T., Taquet, N., Blomquist, G.J., Feyereisen, R., 2012. An insect-specific P450 oxidative decarbonylase for cuticular hydrocarbon biosynthesis. Proceedings of the National Academy of Sciences 109, 14858–14863. 10.1073/PNAS.1208650109

R Core Team, 2014.R: A language and environment for statistical computing.

Rewitz, K.F., O’Connor, M.B., Gilbert, L.I., 2007. Molecular evolution of the insect Halloween family of cytochrome P450s: Phylogeny, gene organization and functional conservation. Insect Biochem. Mol. Biol. 37, 741–753. 10.1016/J.IBMB.2007.02.012

Rinkevich, F.D., Margotta, J.W., Pittman, J.M., Danka, R.G., Tarver, M.R., Ottea, J.A., Healy, K.B., 2015. Genetics, Synergists, and Age Affect Insecticide Sensitivity of the Honey Bee, Apis mellifera. PLoS One 10, e0139841. 10.1371/journal.pone.0139841

Rivera-Colón, A.G., Rehmann, C.T., Kern, A.D., 2026. MKado: a toolkit for McDonald-Kreitman tests of natural selection. bioRxiv. 10.64898/2026.03.02.709122

Rodrigues, C.H.M., Pires, D.E.V., Ascher, D.B., 2020. DynaMut2: Assessing changes in stability and flexibility upon single and multiple point missense mutations. Protein Sci. 30, 60. 10.1002/PRO.3942

Schrödinger LLC, 2015. The PyMOL Molecular Graphics System version 2.0.

Shi, Y., Qu, Q., Wang, C., He, Y., Yang, Y., Wu, Y., 2022. Involvement of CYP2 and mitochondrial clan P450s of Helicoverpa armigera in xenobiotic metabolism. Insect Biochem. Mol. Biol. 140, 103696. 10.1016/j.ibmb.2021.103696

Sim, N.L., Kumar, P., Hu, J., Henikoff, S., Schneider, G., Ng, P.C., 2012. SIFT web server: predicting effects of amino acid substitutions on proteins. Nucleic Acids Res. 40, W452. 10.1093/NAR/GKS539

Suchail, S., Guez, D., Belzunces, L.P., 2000. Characteristics of imidacloprid toxicity in two Apis mellifera subspecies. Environ. Toxicol. Chem. 19, 1901–1905. 10.1002/etc.5620190726

Tajima, F., 1989. Statistical method for testing the neutral mutation hypothesis by DNA polymorphism. Genetics 123, 585–595. 10.1093/genetics/123.3.585

Tarpy, D.R., Vanengelsdorp, D., Pettis, J.S., 2013. Genetic diversity affects colony survivorship in commercial honey bee colonies. Naturwissenschaften 100, 723–728. 10.1007/S00114-013-1065-Y

Tesovnik, T., Zorc, M., Ristanić, M., Glavinić, U., Stevanović, J., Narat, M., Stanimirović, Z., 2020. Exposure of honey bee larvae to thiamethoxam and its interaction with Nosema ceranae infection in adult honey bees. Environmental Pollution 256, 113443. 10.1016/J.ENVPOL.2019.113443

Tosi, S., Sfeir, C., Carnesecchi, E., vanEngelsdorp, D., Chauzat, M.-P., 2022. Lethal, sublethal, and combined effects of pesticides on bees: A meta-analysis and new risk assessment tools. Science of The Total Environment 844, 156857. 10.1016/j.scitotenv.2022.156857

Tsvetkov, N., Bahia, S., Calla, B., Berenbaum, M.R., Zayed, A., 2023. Genetics of tolerance in honeybees to the neonicotinoid clothianidin. iScience 26, 106084. 10.1016/J.ISCI.2023.106084

Vasimuddin, Md., Misra, S., Li, H., Aluru, S., 2019. Efficient Architecture-Aware Acceleration of BWA-MEM for Multicore Systems, in: 2019 IEEE International Parallel and Distributed Processing Symposium (IPDPS). IEEE, pp. 314–324. 10.1109/IPDPS.2019.00041

Volkamer, A., Kuhn, D., Rippmann, F., Rarey, M., 2012. Dogsitescorer: A web server for automatic binding site prediction, analysis and druggability assessment. Bioinformatics 28, 2074–2075. 10.1093/BIOINFORMATICS/BTS310

Volonté, M., Traverso, L., Estivalis, J.M.L., Almeida, F.C., Ons, S., 2022. Comparative analysis of detoxification-related gene superfamilies across five hemipteran species. BMC Genomics 23. 10.1186/s12864-022-08974-y

Wallberg, A., Bunikis, I., Pettersson, O.V., Mosbech, M.-B., Childers, A.K., Evans, J.D., Mikheyev, A.S., Robertson, H.M., Robinson, G.E., Webster, M.T., 2019. A hybrid de novo genome assembly of the honeybee, Apis mellifera, with chromosome-length scaffolds. BMC Genomics 20, 275. 10.1186/s12864-019-5642-0

Wan, N.F., Fu, L., Dainese, M., Kiær, L.P., Hu, Y.Q., Xin, F., Goulson, D., Woodcock, B.A., Vanbergen, A.J., Spurgeon, D.J., Shen, S., Scherber, C., 2025. Pesticides have negative effects on non-target organisms. Nature Communications 2025 16:1 16, 1360-. 10.1038/s41467-025-56732-x

Wieczorek, P., Frąckowiak, P., Obrępalska-Stęplowska, A., 2020. Evaluation of the expression stability of reference genes in Apis mellifera under pyrethroid treatment. Sci. Rep. 10, 1–15. 10.1038/S41598-020-73125-W

Wragg, D., Eynard, S.E., Basso, B., Canale-Tabet, K., Labarthe, E., Bouchez, O., Bienefeld, K., Bieńkowska, M., Costa, C., Gregorc, A., Kryger, P., Parejo, M., Pinto, M.A., Bidanel, J.P., Servin, B., Le Conte, Y., Vignal, A., 2022. Complex population structure and haplotype patterns in the Western European honey bee from sequencing a large panel of haploid drones. Mol. Ecol. Resour. 22, 3068–3086. 10.1111/1755-0998.13665

Wu, H., Li, Z., Zhong, Z., Guo, Y., He, L., Xu, X., Mao, Y., Tang, D., Zhang, W., Jin, F., Pang, R., 2025. Insect Cytochrome P450 Database: An Integrated Resource of Genetic Diversity, Evolution and Function. Mol. Ecol. Resour. 25, e14070. 10.1111/1755-0998.14070

Wu, M.C., Chang, Y.W., Lu, K.H., Yang, E.C., 2017. Gene expression changes in honey bees induced by sublethal imidacloprid exposure during the larval stage. Insect Biochem. Mol. Biol. 88, 12–20. 10.1016/J.IBMB.2017.06.016

Wu, Y., Zheng, Y., Li-Byarlay, H., Shi, Y., Wang, S., Zheng, H., Hu, F., 2020. CYP6AS8, a cytochrome P450, is associated with the 10-HDA biosynthesis in honey bee (Apis mellifera) workers. Apidologie 51, 1202–1212. 10.1007/S13592-019-00709-5

Wu, Y.Q., Zheng, H.Q., Corona, M., Pirk, C., Meng, F., Zheng, Y.F., Hu, F.L., 2017. Comparative transcriptome analysis on the synthesis pathway of honey bee (Apis mellifera) mandibular gland secretions. Sci. Rep. 7, 1–10. 10.1038/S41598-017-04879-Z

Xiao, T., Lu, K., 2022. Functional characterization of CYP6AE subfamily P450s associated with pyrethroid detoxification in Spodoptera litura. Int. J. Biol. Macromol. 219, 452–462. 10.1016/J.IJBIOMAC.2022.08.014

Zanger, U.M., Schwab, M., 2013. Cytochrome P450 enzymes in drug metabolism: regulation of gene expression, enzyme activities, and impact of genetic variation. Pharmacol. Ther. 138, 103–141. 10.1016/J.PHARMTHERA.2012.12.007

Zhou, S.F., Liu, J.P., Chowbay, B., 2009. Polymorphism of human cytochrome P450 enzymes and its clinical impact. Drug Metab. Rev. 41, 89–295. 10.1080/03602530902843483

Zou, F., Guo, Q., Shen, B., Zhu, C., 2019. A cluster of CYP6 gene family associated with the major quantitative trait locus is responsible for the pyrethroid resistance in Culex pipiens pallen. Insect Mol. Biol. 28, 528–536. 10.1111/IMB.12571

